# AI-BioMech: Deep Learning Prediction of Mechanical Behavior in Aperiodic Biological Cellular Materials

**DOI:** 10.64898/2026.02.24.707699

**Authors:** Haleema Sadia, Marcelo A. Dias, Parvez Alam

## Abstract

We introduce AI-BioMech, a deep learning based framework that directly predicts the mechanical response of cellular structures from 2D images, eliminating the need for manual geometry definition and traditional finite element simulations. The framework is trained on synthetic datasets representing biological cellular structures and benchmarked against real experimental data. Finite element analysis (FEA) based labeling is used to generate pixel level annotations for semantic segmentation, enabling accurate identification of stress and strain distributions. By learning spatial and hierarchical patterns from these annotations, the model automatically extracts complex features to predict cellular material responses under compressive loading conditions. Transfer learning with fine tuning by using the DeepLabv3 architecture with ResNet50, ResNet101, and Inception ResNetV2 backbones enhances prediction accuracy and generalization from limited datasets. Model predictions are validated against experimental results and Digital Image Correlation (DIC) measurements, demonstrating strong agreement with physical observations. The results show that AI-BioMech achieves up to 99% prediction accuracy while significantly outperforming traditional methods in computational speed and scalability.

## 1 Introduction

An ability to predict the mechanical behavior of cellular materials has broad applicability across various fields [1]. The modeling of high resolution intricate microstructures is nevertheless not always trivial, often demanding considerable computational resources, limiting both large scale simulations and any extensive exploration of structure-properties design spaces [2, 3]. The highly irregular and complex geometries of biological structures makes manual modeling and conventional simulation approaches, such as finite element analysis (FEA), time consuming and error prone [4] [5], motivating the need for efficient, and automated approaches.

Advances in artificial intelligence (AI) offer promising solutions to these challenges. Microstructure analysis using AI and image processing has become an active area of research [6–10], with convolutional neural networks (CNNs) demonstrating strong predictive capabilities. CNNs automatically extract hierarchical features from raw image data, enabling the identification of microstructural patterns such as pore geometry, cell wall thickness, and textures. These learned features support the accurate prediction of material properties, while enabling the automated detection, quantification, and classification of microstructural phases and defects [11]. By leveraging AI algorithms, it may therefore be possible to overcome some of the limitations of traditional simulation methods, while concurrently enabling the capture of complex material behavior [12].

When comparing the different AI approaches, semantic segmentation proves itself to be a powerful technique in predicting material properties at the pixel level [13]. By assigning a class label to each pixel, semantic segmentation enables detailed analysis of microstructural features, including material phases [7], structural defects [14], and classes such as metals, polymers, ceramics, and glass [6, 15]. Architectures like U-Net capture fine-scale details through encoder–decoder structures, while DeepLab leverages atrous convolutions to model multi-scale features [6,8,9]. Other models, including FCN, SegNet, and PSPNet, have also been applied to segment small and irregular geometries. The use of pre-trained CNN backbones such as ResNet, VGG, and EfficientNet enhances feature extraction through transfer learning, improving segmentation accuracy and enabling the precise prediction of material properties [16]. While these approaches have demonstrated effectiveness in phase identification [17, 18], their potential for predicting spatially resolved mechanical fields, such as stress–strain distributions in cellular materials, remains underexplored [19].

Machine learning (ML) more broadly has also been applied to material property prediction, including Support Vector Machines (SVM) for micro-CT segmentation, Random Forests (RF) for concrete property prediction, and gradient boosting methods for small datasets [20–22]. Advanced ML approaches, such as Bi-LSTM, GANs, and pixel-based LSTM frameworks, have enabled the rapid prediction of dynamic responses, optimized lattice designs, and enhanced additive manufacturing performance [23–29]. Despite these advances, significant challenges persist for cellular materials, including the difficulty of linking complex microstructures to mechanical behavior, limited high-quality datasets, resource intensive training of deep learning models, limited generalization across materials, and the “black-box” nature of many models [30, 31]. These challenges elucidate a need for integrated AI frameworks that can efficiently predict stress-strain distributions and mechanical responses in complex cellular structures.

Transfer learning (TL) is a powerful method that can address the challenges associated with limited dat availability. TL adapts models pretrained on large datasets, such as ImageNet, to smaller, specialized datasets, reducing the need for extensive labeled data and computational resources [31–34]. Fine-tuning pretrained models on cellular material images improves accuracy and efficiency in predicting mechanical behavior. Prior studies demonstrate the effectiveness of TL in material science, such as predicting steel corrosion in concrete [32], describing stress–strain responses in elastoplastic solids with RNNs, and classifying construction material conditions using Inception V3 [33]. For cellular materials, TL enables learning from microstructural images with limited experimental or FEA generated data, facilitating the accurate prediction of stress–strain data.

While deep learning and transfer learning offer significant advantages over conventional machine learning approaches in material properties predictions, their effectiveness depends on the availability of large, high quality, labeled datasets [35]. For cellular materials, generating such labels is particularly challenging. Acquiring labels through FEA simulations is time consuming, and manually modeling complex, aperiodic, and irregular cellular structures is challenging. The scarcity of labeled data in the target domain limits a model’s ability to generalize, reducing its accuracy in predicting mechanical and functional properties. Additional challenges include domain gaps between source and target datasets and the complexity of fine-tuning models for specific material properties. In aperiodic cellular structures, small datasets can cause overfitting, limiting generalization. Addressing these issues requires careful model selection, data alignment, and optimization [36, 37], highlighting the need for automated frameworks that efficiently generate labels and predict mechanical behavior in biological cellular structures.

In consideration of the above mentioned challenges, we introduce AI-BioMech, a 2D image-based deep learning framework, developed to predict the mechanical behavior of complex aperiodic cellular materials, while providing a fast and scalable alternative to FEA. To overcome limitations in accessing labeled data, synthetic datasets are generated by defining cellular geometries, assigning material properties, and solving partial differential equations (PDEs) to obtain stress and displacement fields, which serve as training targets. The framework integrates semantic segmentation and transfer learning. Semantic segmentation enables the pixel-level identification of structural regions and material behavior (stress-strain distribution), while transfer learning with pre-trained networks (DeepLab with ResNet50, ResNet101, and Inception-ResNetV2) captures fine and large scale patterns, improving accuracy, generalization, and efficiency. By combining FEA-based labeling alongside deep learning, AI-BioMech addresses modeling challenges associated with irregular geometries, data scarcity, and computational cost, enabling rapid, highly accurate stress–strain predictions, and supporting the exploration of 2D design space.

## 2 Methodology

This research is inspired by biological cellular structures such as sponge, wood, and trabecullar bone, each of which exhibits highly irregular geometries with diverse pore shapes, sizes, and cell wall thicknesses. Here, we aim to predict the mechanical behavior of complex, aperiodic cellular architectures by combining synthetic data generation, image-based finite element analysis (FEA), and deep learning. Our outputs will include predicted stress–strain distributions within irregular cellular features. Figure 1 illustrates the overall framework of the proposed study.

**Figure 1.**
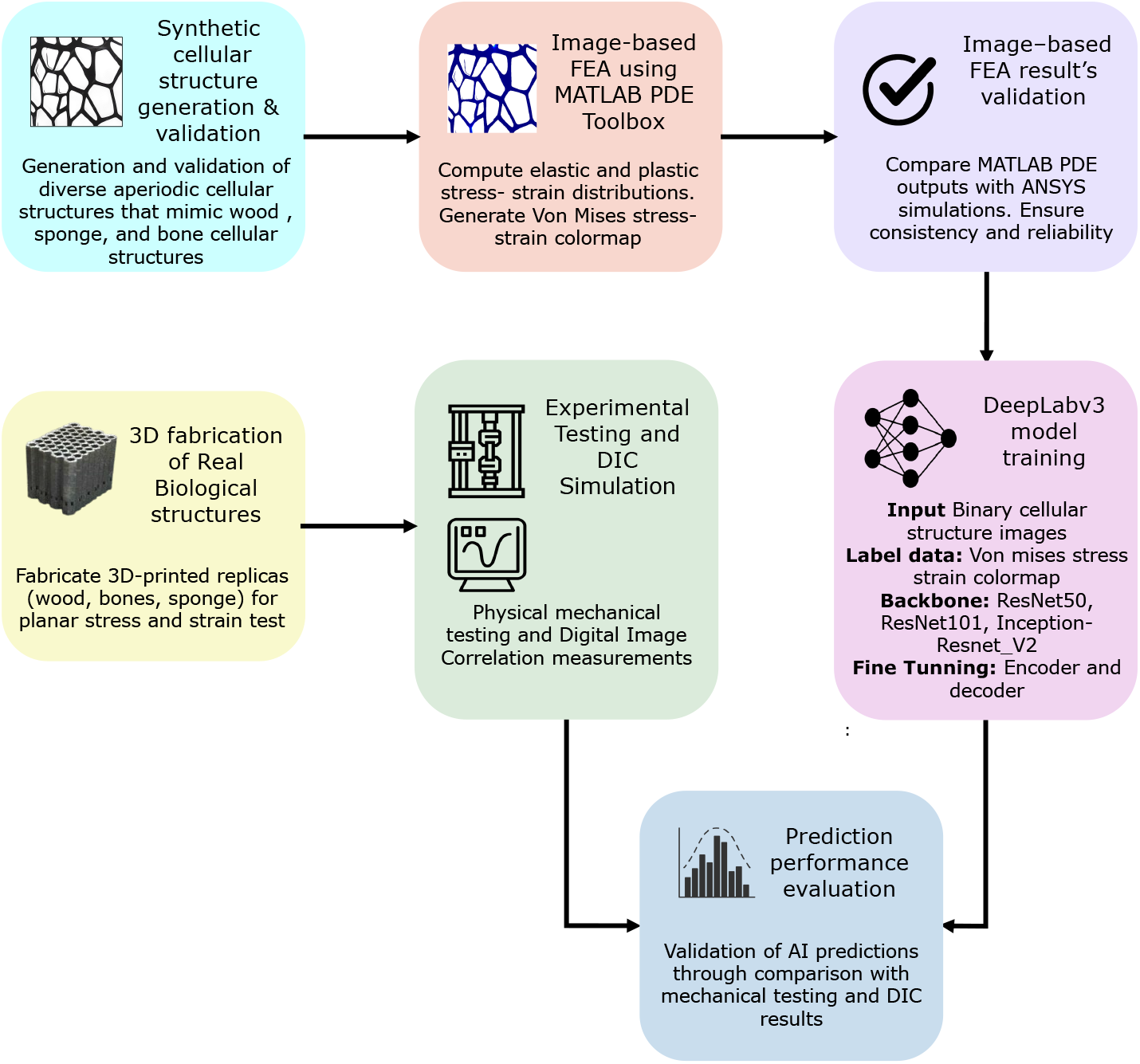
Overview of the end-to-end AI-BioMech framework: Synthetic dataset of complex cellular structures are first generated to replicate real biological geometries and benchmarked against real biological cellular datasets. Image-based FEA is then performed to compute stress and strain distributions, which are validated against ANSYS (Ansys Inc., Canonsburg, PA, USA) simulations. The validated stress–strain maps serve as ground-truth labels to train a DeepLabv3 model for pixel-wise prediction of mechanical fields. After achieving optimal training performance, the model is tested on 3D-printed biological exemplars including: wood, bone, and sponge, subjected to static compressive loading. Experimental stress–strain data obtained via Digital Image Correlation (DIC) and mechanical testing are compared against the model predictions, providing a rigorous means to validating of the AI-BioMech framework

To train the model, we first generated a comprehensive synthetic dataset (binary images) of cellular geometries designed to mimic the structural diversity and irregularity of real biological morphologies. To ensure the synthetic dataset accurately represents real-world cellular structures, we validated it by performing correlation analyses between synthetic and real biological cellular dataset features, confirming that the synthetic geometries and their mechanical responses closely mimic real-world features. After validation, each synthetic structure was analyzed to perform image-based FEA, evaluating both elastic and plastic stress–strain responses. The resulting displacement and stress-strain distributions were converted into high resolution colormaps, which served as pixel-level labels for supervised learning.

To ensure the accuracy of the image-based FEA, the results were validated against ANSYS simulations, confirming consistency with standard commercial FEA outputs. Following validation, the synthetic cellular structures and their corresponding stress–strain colormaps were used to train a DeepLabv3 semantic segmentation network. Three backbone encoders, ResNet-50, ResNet-101, and Inception-ResNetV2, along with encoder-decoder components including the Atrous Spatial Pyramid Pooling (ASPP) module, were fine-tuned to map binary cellular inputs to their corresponding stress–strain outputs. After achieving optimal training performance, the model was tested on completely unseen real biological cellular structures under both plane stress and plane strain conditions. For this purpose, 3D-printed replicas of biological samples, including wood, and sponge, were fabricated as physical test specimens. The 2D stress–strain predictions were compared against experimental data (compression tests) and Digital Image Correlation (DIC). This comparison allowed us to evaluate the model’s accuracy and its ability to generalize from synthetic training data to real-world biological structures.

Overall, this methodology combines synthetic dataset generation, physics-based FEA labeling, semantic segmentation, transfer learning, and experimental validation to establish a robust AI-BioMech framework for predicting the mechanical behavior of complex, irregular biological cellular materials directly from images. This approach eliminates the need for manual geometry definition and traditional FEA frameworks, while enabling accurate stress–strain estimation and supporting efficient design exploration and optimization of cellular structures. The individual parts of this framework will now be covered in greater detail in the sections that follow.

### 2.1 Generation of synthetic datasets for biological cellular structures

Synthetic datasets were created initially, by modeling regular cellular structures, including honeycomb hexagonal, square, and triangular lattices. When natural cellular materials such as wood, bone, and sponge, are projected in 2D, they exhibit stochastic patterns that follow Voronoi honeycomb [38–44]. As such, we introduced controlled perturbations to hexagonal, square and triangle honeycomb lattices, to replicate these natural (Voronoi-like) irregularities, ensuring the synthetic dataset captures the structural diversity and complexity of real-world cellular materials.

Each lattice type was defined using a representative unit cell characterized by its side lengths and interior angles, as illustrated in Figure 2. Specifically, a hexagonal lattice unit cell consists of three sides (*l*_1_, *l*_2_, *l*_3_) and three angles (*α, β, γ*); a square lattice has four sides (*l*_1_–*l*_4_) and four angles (*α*–*δ*); and a triangular lattice comprises six sides (*l*_1_–*l*_6_) and six angles (*α, β*_1_, *β*_2_, *δ*_1_, *δ*_2_, *γ*). In a regular honeycomb lattice, all sides are equal and the interior angles sum to 360^°^.

**Figure 2.**
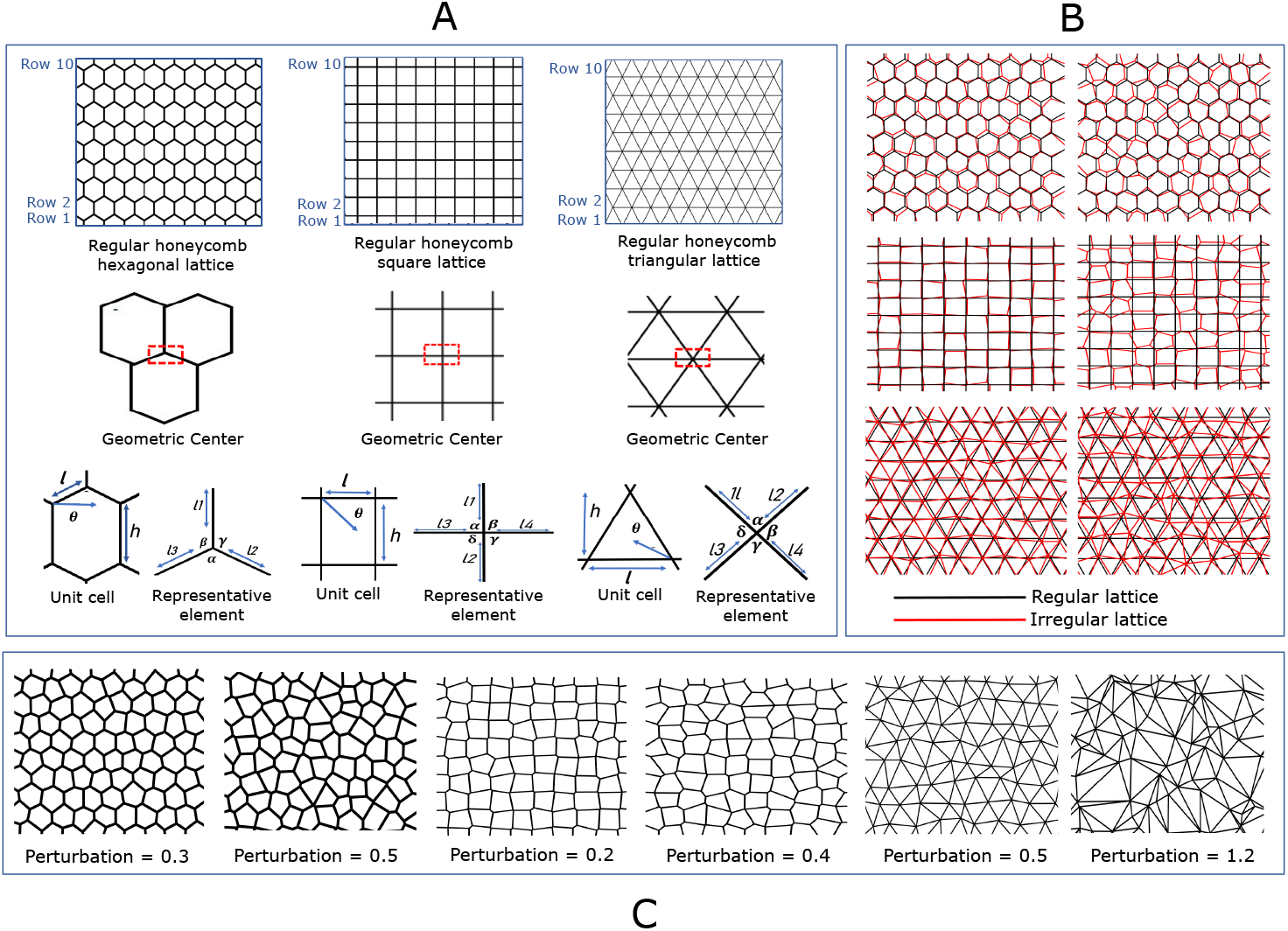
(A) Illustration of hexagonal, square, and triangular honeycomb lattices with outlined unit cells showing height and width dimensions. Representative elements are marked within each lattice, replicate to form the entire structure: the hexagonal element has three sides (*l*_1_, *l*_2_, *l*_3_) and angles (*α, β, γ*) summing to 360^°^; the square element has four sides (*l*_1_–*l*_4_) and angles (*α*–*δ*) summing to 360^°^; and the triangular element has six sides (*l*_1_–*l*_6_) and angles (*α, β*_1_, *β*_2_, *δ*_1_, *δ*_2_, *γ*) summing to 360^°^. (B) Illustration of synthetic cellular structures generated by introducing geometric perturbations into the regular lattices. The perturbations alter side lengths and angles, producing irregular, non-periodic patterns that capture structural variability and complexity, while preserving overall lattice integrity. (C) Examples illustrate the range of irregularities achievable through controlled perturbations, which form the basis for generating synthetic datasets to train the AI-BioMech framework.

To better mimic the Voronoi type irregularities observed in 2D projections of real biological cellular structures, controlled geometric perturbations were applied to the regular honeycomb lattices. Using hexagonal unit cells as an example: both the side lengths and interior angles of each hexagonal unit cell were treated as independent variables, while geometric integrity was maintained through constraints such as shared edge consistency and the preservation of the sum of interior angles. Under these constraints, the perturbed angles must satisfy the condition that their sum remains equal to 360^°^, as shown in Equation 1:

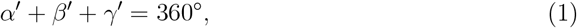

where, *α*^*′*^, *β*^*′*^, and *γ*^*′*^ are peturbed angle, the relationship between the original angles *α, β*, and *γ* and the perturbed is defined through the angular perturbations *ϵ*_*α*_, *ϵ*_*β*_, and *ϵ*_*γ*_ as shown in Equation 2:

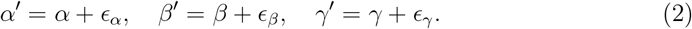

By substituting these perturbed angle expressions into the angular sum constraint, Equation 1 becomes Equation 3:

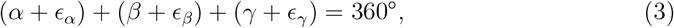

which ensures preservation of the overall hexagonal geometry, as these perturbations satisfy *ϵ*_*α*_+*ϵ*_*β*_ +*ϵ*_*γ*_ = 0. In this study, the perturbation parameter is defined as *ϵ* ∈ [−2, +2] with a step size of 0.1, indicating that the angular perturbations *ϵ*_*α*_, *ϵ*_*β*_, and *ϵ*_*γ*_ for each hexagonal cell lie within this interval. A similar perturbation scheme was applied to the square and triangular lattices, ensuring variations in both angles and side-lengths were defined to preserve the geometrical integrity of the unit cell.

After generating perturbed unit cells, we assembled them into multicellular configurations to progressively increase structural complexity within the dataset. We started with a single cell image and systematically expand to 2×2 (4-cell), 3×3 (9-cell), 4×4 (16-cell), 5×5 (25-cell), and 6×6 (36-cell) arrangements. This approach produces irregular cellular patterns at each stage, resulting in a total of 80,000 images for training, with each stage introducing higher levels of geometric variability and complexity.

#### 3.1.1 Data augmentation

To further enrich the dataset, data augmentation techniques were applied as shown in Figure 3. To simulate natural variability in cell wall thickness, random dilation and erosion operations were applied to different regions of the binary images. Each region undergoes stochastic morphological modification, with the magnitude of dilation or erosion selected from a specified thickness range of [5, 50] pixels. Dilation expands cell boundaries, increasing wall thickness, while erosion contracts them, reducing thickness. Applying these operations randomly across the images introduces realistic heterogeneity, closely reflecting the variations found in real biological structures. This stochastic modification increases geometric realism and helps the model generalize to unseen cellular patterns.

**Figure 3.**
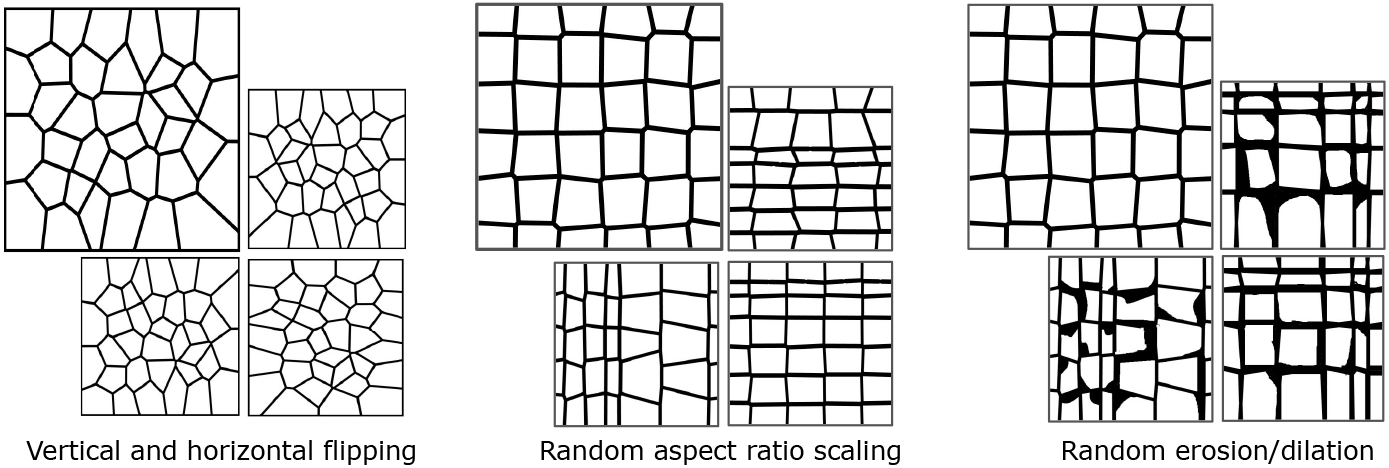
Illustration of the data augmentation techniques applied to the dataset to enhance diversity. These techniques include vertical and horizontal flipping, random aspect ratio scaling from 1× to 20×, and random dilation and erosion applied within a range of [5, 50]. By applying these transformations, the model is exposed to a wider range of variations, improving its ability to generalize to different orientations, positions, and structural scales in real world scenarios

Additional dataset enrichment was then achieved through the application of data augmentation techniques including: random rotations within 0^°^–360^°^, vertical and horizontal flipping, and random aspect ratio scaling, where the width and height of cells were independently varied. Combined with the stochastic dilation and erosion operations, these augmentations introduce natural variations in cell shape, elongation, and anisotropy while maintaining structural realism. These methods enhanced the diversity of our dataset, helping the model learn robust, orientation-invariant features, and better handle irregular structures. Following these augmentations, the final dataset expands to approximately 350,000 images.

#### 2.1.2 Real world benchmarking of the synthetic dataset

Following data augmentation, the synthetic dataset was validated against real world biological cellular structures including wood, sponge, and bone to ensure that the key structural features of real samples are accurately captured. A feature comparison framework based on Convolutional Neural Networks (CNNs) was employed [45,46]. Specifically, ResNet-101 was used to extract and compare deep feature representations from a large-scale synthetic dataset comprising 350,000 images and a real-world dataset consisting of 500 images. The real-world dataset was assembled from publicly available images of biological cellular structures sourced from open-access platforms such as figshare and Wikimedia Commons, including images released under Creative Commons (CC 4.0) licenses.

Since our synthetic dataset consists of binary images, all real world biological images were first converted to binary representations and subsequently mapped to grayscale, ensuring that both synthetic and real images lie on the same representational plane and encode only geometric information. All images were then resized to a uniform resolution of 224 × 224 to match the network input layer, enabling consistent geometry based feature comparison. We employed the ResNet-101 architecture, chosen for its residual connections that effectively capture both fine grained local textures and high level global patterns. Its deep architecture enables enhanced feature extraction, making it suitable for the analysis of complex biological structures [47].

As shown in Figure 4, each image (synthetic and real) was passed through the network, and feature vectors were extracted from multiple layers to capture multi-scale information: intermediate convolutional layers for local edge and texture patterns, and the final global average pooling layer (pool5) for global structural representation. All feature vectors were L2-normalized to unit length to ensure that comparisons focus on angular alignment as well as absolute magnitude. The structural similarity between synthetic and real images was then quantified using cosine similarity and Euclidean distance, providing a rigorous measure of how well the synthetic dataset captures the key features present in real biological samples.

**Figure 4.**
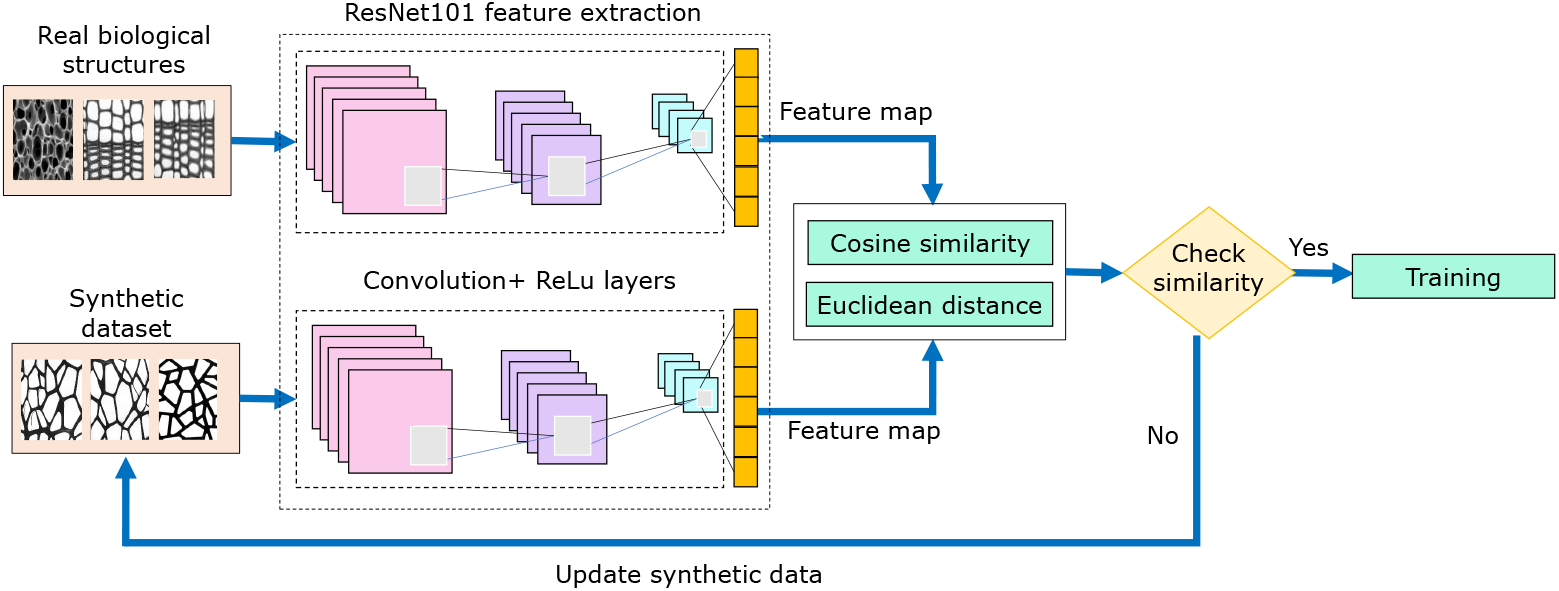
Workflow for validating the synthetic dataset against real world biological structures. Real images (500) and synthetic images (350,000) are processed using a ResNet-101 network for feature extraction. The extracted features from both datasets are compared using cosine similarity and Euclidean distance to assess structural similarity. The synthetic dataset is iteratively refined until the feature maps achieve the desired similarity, after which it is finalized for model training.

The cosine similarity between two feature vectors **f**_real_ and **f**_syn_ is defined by Equation 4:

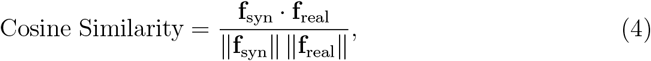

where, **f**_real_, and **f**_syn_ are the normalized feature vectors of real and synthetic images, respectively. A value of 1 indicates identical angular alignment, whereas 0 indicates orthogonal features. Using this procedure, each synthetic image was compared against all 500 reference real images, producing a vector of 500 similarity scores per synthetic image. For the entire synthetic dataset of 350,000 images, this results in a total of 350,000 × 200 cosine similarity values, which were subsequently analyzed using histograms and tail distributions to assess the structural alignment and similarity of synthetic images relative to the reference dataset.

Euclidean distance provides a measure of the absolute difference between the features computed, as shown by Equation 5:

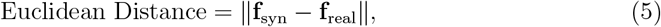

where the Euclidean distance of 0 indicates that the feature vectors are identical, i.e. there is no difference between the corresponding features of the two images. Higher Euclidean distance values reflect greater dissimilarity in the feature space. This methodology provided a robust, multi-level validation, demonstrating that the synthetic dataset faithfully reproduced the structural characteristics of real biological samples despite the large imbalance between synthetic and real data [48, 49]. The synthetic dataset was iteratively updated until it satisfied the similarity criteria, after which it was used for training.

### 2.2 Pixel-based labeling using FEA-generated color maps

Pixel-level labels were generated from image-based Finite Element Analysis (FEA) color maps that were used to train the DCNN model. First, each binary image was preprocessed to extract the black regions, which were defined as the regions of interest (ROI) for meshing, as illustrated in Figure 5. Within these ROIs, a triangular mesh was generated using a uniform grid spacing of 0.01, ensuring consistent discretization for analysis. The computational geometry was then constructed by assigning the appropriate properties for the material (e.g., bone, wood, etc.) and geometrical dimensions to accurately represent the physical structure.

**Figure 5.**
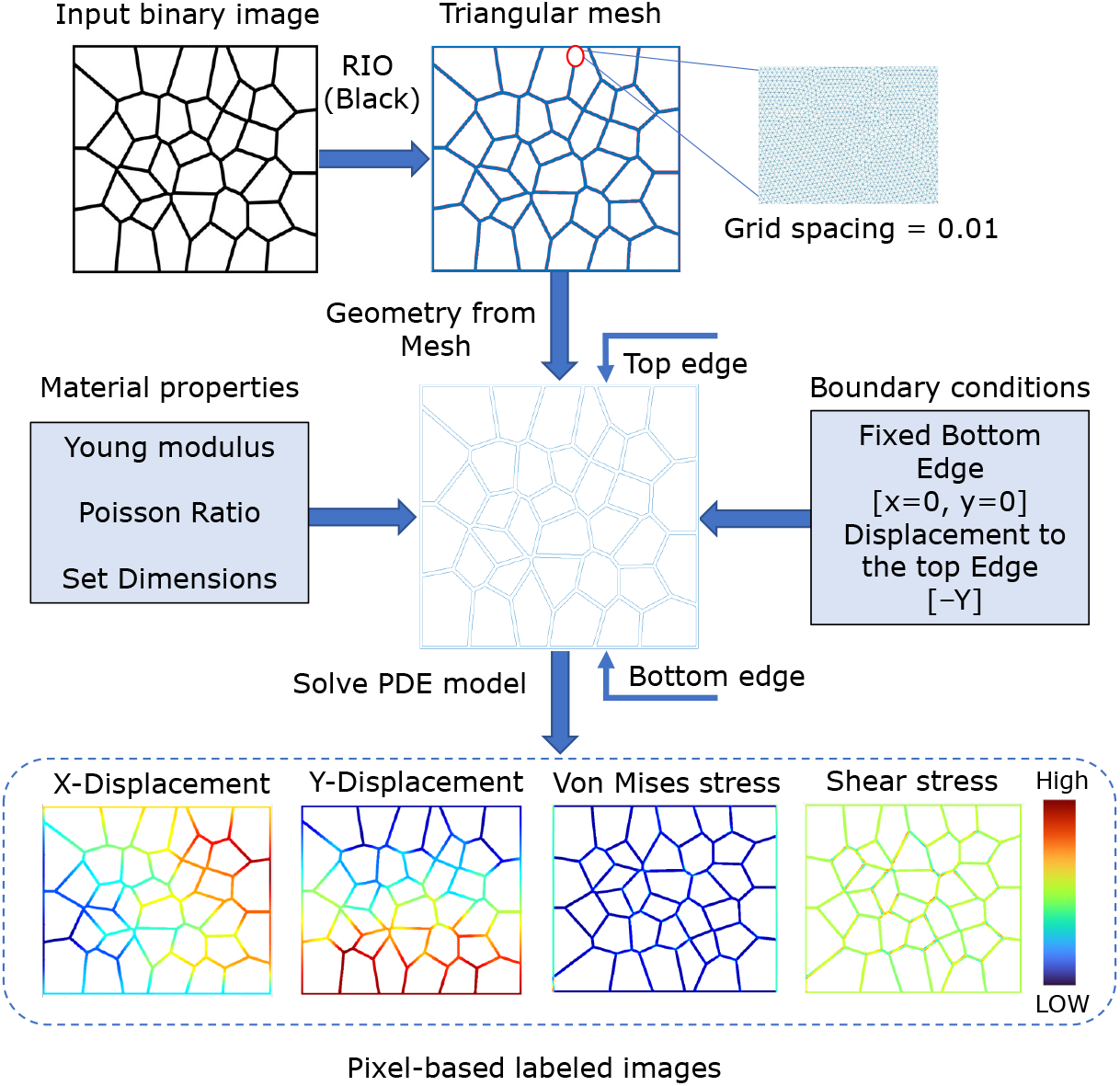
Illustration of the pixel-based labeling workflow used to generate ground-truth data from image-based FEA simulations. The process begins by extracting the region of interest (ROI) from a binary image and generating a triangular mesh with uniform spacing. Material properties and boundary conditions are applied, with the bottom edges fixed and a variable downward displacement imposed on the top edges. FEA simulations produce displacement fields (X and Y) and stress fields (von Mises and shear), which are converted into Jet64 colormap images, where each pixel encodes the corresponding local stress or displacement value. These labeled images form the training dataset for the DCNN model, enabling it to learn spatial stress patterns and predict stress distributions in complex, aperiodic cellular structures

Boundary conditions were imposed by fixing the bottom edges and applying a displacement vector of [0, −*Y*] to the top edges, with the displacement magnitude incrementally varied from 0.1% to 10% of the total structural length. The model was then solved numerically to evaluate key mechanical responses, including X- and Y-displacements, von Mises stress, and normal and shear stresses. These simulations produced detailed stress and displacement maps, which were subsequently converted into pixel-based labels using the Jet64 colormap, where blue represents low stress and red represents high stress.The accuracy of the image-based FEA results was validated by comparison against equivalent ANSYS simulations, ensuring both consistency and reliability of the generated stress and displacement fields.

This framework not only enables the identification of stress concentrations but also captures the spatial variations of stress across highly irregular and aperiodic cellular architectures. By integrating FEA-generated datasets with deep learning, the approach provides a powerful and robust tool for analyzing and understanding stress–strain behavior in complex, aperiodic structures, facilitating accurate prediction and insight into the mechanical performance of diverse cellular materials [50].

### 2.3 Transfer learning

Following dataset preparation and labeling, we employed transfer learning to enhance the performance of our semantic segmentation task using the DeepLabv3 architecture with three backbone variations: ResNet50, ResNet101, and Inception-ResNetV2. These pre-trained CNN backbones provide a strong starting point by leveraging features learned from large-scale datasets, allowing the model to generalize effectively on our FEA-labeled dataset. Fine-tuning and hyperparameter optimization were performed for all three backbones to identify the best performing model. The optimal backbone was then selected based on its accuracy and segmentation quality, ensuring the robust prediction of stress and strain distributions in cellular structures.

A key component enabling this performance is convolutional feature extraction, which allows CNNs to identify meaningful spatial patterns from the input images. Mathematically, this involves applying a convolution operation using a kernel, a smaller *K*^*′*^ *× K*^*′*^ matrix of weights, on the input image *I*, represented as a *H*^*′*^ *× W*^*′*^ matrix, where *H* and *W* denote the height and width of the image, respectively. The kernel slides across the input with a specified stride *s*, performing element-wise multiplication and summation at each position. The resulting output feature map *O*, of size *H*^*′*^ *× W*^*′*^, captures localized patterns essential for accurate stress and strain prediction, as computed by Equation 6:

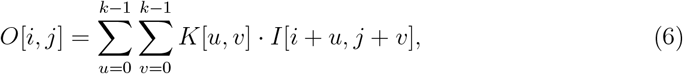

where *K*[*u, v*] represents the kernel weights, and *I*[*i* + *u, j* + *v*] is the corresponding input value under the kernel, where *i* = 0, …, *H*^*′*^−1 and *j* = 0, …, *W*^*′*^−1 index the vertical and horizontal positions of the output feature map *O*, and *u* = 0, …, *k*−1 and *v* = 0, …, *k*−1 index the vertical and horizontal positions of the kernel *K*. The stride determines the step size for the kernel movement. Zero-padding of size *p* can be added to the input feature map to preserve spatial dimensions, where *p* denotes the number of padding pixels added to each side of the input. The kernel size is denoted by *k*, representing the height and width of the convolutional filter. The resulting output dimensions *H*^*′*^ and *W*^*′*^ are shown by Equations 7 and 8, respectively:

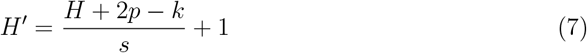

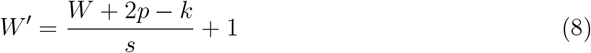

Following the convolution operation, a non-linear activation function such as ReLU is applied element-wise to enhance feature representation, as defined in accordance with Equation 9:

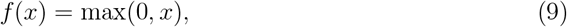

where *x* represents a scalar element of the convolution output feature map before activation. Following feature extraction through convolution, max pooling is often applied to reduce the spatial dimensions and retain only the most prominent features. Max pooling divides the feature map into non-overlapping regions of size *p × p* and selects the maximum value from each region. For a feature map *F*, the pooled output *P* is given by Equation 10:

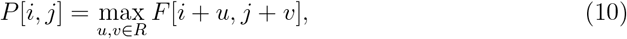

where *R* represents the region covered by the pooling window.

To stabilize the training process and improve convergence, batch normalization was applied after convolution or activation layers. Batch normalization normalizes the output of a layer by adjusting and scaling the activations, ensuring that they have a mean of 0 and a variance of 1. For a batch of activations *x*, the normalized output 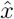 is given by Equation 11:

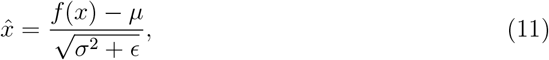

where, *µ* and *σ*^2^ are the mean and variance of the batch, and *ϵ* is a small constant to prevent division by zero. The normalized values are then scaled and shifted using learnable parameters *γ* and *β*, producing the final output *y* of the batch normalization layer, as shown in Equation 12:

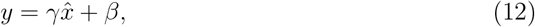

where *y* represents the transformed activation value that is forwarded to the subsequent layer.

After feature extraction by the backbone network, the resulting feature map was passed to the Atrous Spatial Pyramid Pooling (ASPP) module, which enhances multi-scale contextual understanding as shown in Figure 6. The ASPP applies atrous convolutions with varying dilation rates (1, 3, 6, 12, and 18) to expand the receptive field without losing spatial resolution, enabling the network to capture both fine-grained details and long-range context. Global average pooling was also applied to incorporate global contextual information by pooling the entire feature map into a single value. The outputs of the atrous convolutions and global pooling are concatenated and refined using a 1 × 1 convolution, producing an enriched feature map. This multi-scale encoder was then forwarded to the decoder, allowing the model to accurately segment stress–strain regions of varying scales within complex cellular structures.

**Figure 6.**
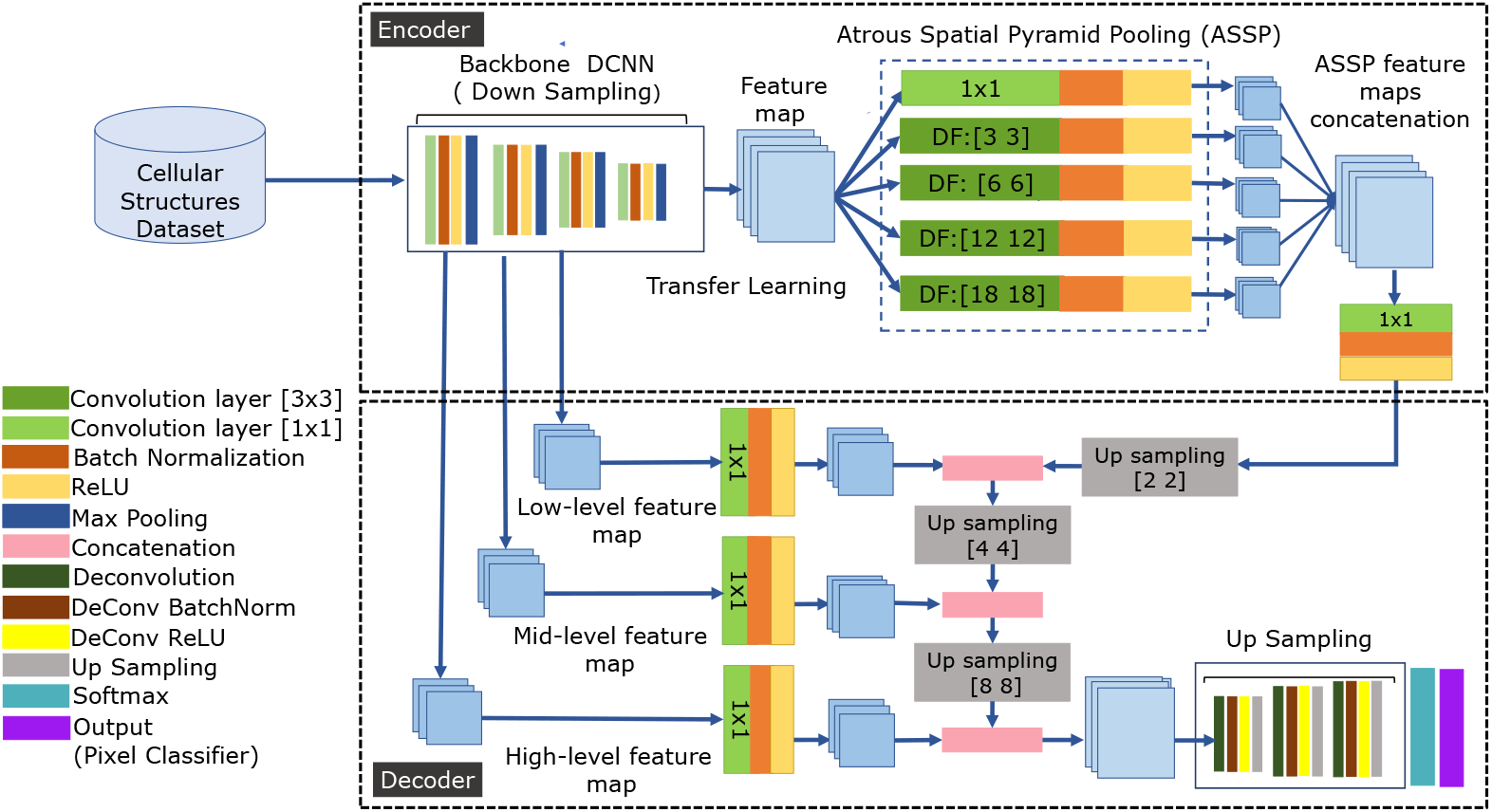
Detailed illustration of the network architecture comprising an encoder-decoder structure. The encoder extracts features from the input data through multiple layers of convolutional operations, progressively reducing spatial dimensions while retaining important information. After feature extraction, an ASSP (Atrous Spatial Pyramid Pooling) module is applied to capture multi-scale contextual information. These concatenated feature maps are then passed through an upsampler before being sent for further upsampling and refinement in the decoder for accurate predictions and pixel classification.

In our proposed method, the decoder progressively reconstructs the spatial resolution of the feature maps while integrating multi-level contextual information. The feature map from the ASPP module was first upsampled by a 2 × 2 transpose convolution and concatenated with high-level encoder features to preserve fine details. This was followed by a 4 × 4 upsampling and concatenation with middle-level features, capturing additional contextual information. Finally, an 8 × 8 upsampling integrates low-level features, enhancing spatial resolution and fine-grained detail. By fusing low, middle, and high-level features at each stage, the decoder combines local details with global context, improving segmentation accuracy for complex cellular structures. The fused feature maps were further refined using an upsampling layer, followed by a softmax for pixel-wise classification, and post-processed with deconvolution, ReLU activation (DeReLU), and batch normalization (DeBatchNormalization) to ensure high-resolution, stable, and precise segmentation predictions.

To train the semantic segmentation model, key hyperparameters were carefully selected to encourage efficient learning and robust performance. The optimizer was evaluated using Adam, SGD, and SGD with momentum, to determine the most effective update strategy. Learning rates of 0.0001, 0.001, and 0.01 were tested to balance convergence speed and stability. Batch sizes of 16, 32, and 64 were explored to optimize memory usage while maintaining stable training. The model was trained for 100 epochs with early stopping to prevent overfitting when validation performance plateaued. A learning rate scheduler was implemented to reduce the learning rate progressively during training, allowing finer adjustments in later stages. Categorical cross-entropy loss was used to measure prediction errors, thereby guiding the optimizer. Dropout (rate 0.3-0.5) was applied after the convolutional layering to improve generalization by randomly deactivating neurons. Weights were initialized using He initialization to ensure appropriate starting values. ReLU activation was used throughout the network to introduce non-linearity, while softmax activation at the output layer provided pixel-wise classification probabilities.

All fine-tuning and hyperparameters were iteratively optimized using the validation dataset, continuing until an optimal combination was identified that ensured both accurate and stable model training. The finalized model was then deployed within the AI-BioMech framework, enabling reliable and efficient prediction of mechanical behavior directly from images of cellular structures.

### 2.4 Post-processing for physically interpretable predictions

Once the model achieved satisfactory segmentation accuracy, the predicted displacement color maps were combined with material properties and constitutive relations to compute the corresponding numerical stress and strain fields. This post-processing step enables a physically meaningful interpretation of the network outputs. Although the proposed deep learning framework is not physics-informed, it is able to produce accurate and mechanically consistent predictions by integrating semantic segmentation results with material properties after inference.

While physics-informed neural networks (PINNs) embed governing equations directly into the training process, their application becomes challenging for highly irregular or aperiodic cellular structures [51]. PINNs typically require explicit definitions of governing equations and boundary conditions over the entire computational domain, leading to increased computational cost and implementation complexity for non-uniform geometries [52–54]. In contrast, our approach makes use of DeepLabv3-based semantic segmentation to learn spatial displacement and stress patterns directly from FEA-generated data. By decoupling physics from the learning phase, the network efficiently captures complex local features across irregular structures. Material properties are subsequently incorporated during post-processing to reconstruct physically meaningful stress and strain fields. This strategy avoids the convergence difficulties and high computational overhead associated with training PINNs from scratch, while maintaining quantitatively accurate predictions for complex cellular geometries.

AI-BioMech predicts the deformation of highly aperiodic cellular structures in the form of color maps. Each pixel in the predicted image represents a scalar quantity, encoded using a Jet64 colormap, where blue represents the minimum value (0), red represents the maximum value (1), and intermediate values are represented by shades of green and yellow. This colormap provides a visual representation of the displacement and stress distributions across the structure. **C**(*X, Y*) denotes the RGB value at pixel (*X, Y*) as in Equation 13:

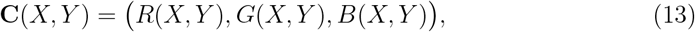

where RGB values are first normalized to the range [0,1] as given in Equation 14.

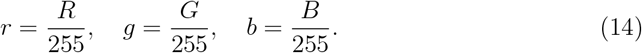

With Jet64, each normalized color corresponds to a discrete color index *k* ∈ [0, 63]. The normalized scalar value *S*(*X, Y*) is computed as Equation 15:

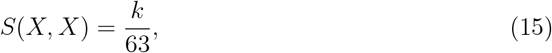

where *S*(*X, Y*) ∈ [0, 1] represents the relative magnitude of the predicted quantity.

The network predicts the displacement components along the *X*- and *Y* -directions as Equation 16:

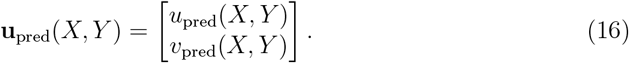

The prediction is normalized, the physical displacement field is reconstructed from the applied boundary conditions. For example, with a prescribed vertical displacement Δ*Y* at the top edge and a fixed bottom edge, the average axial strain along *Y* is described by Equation 17:

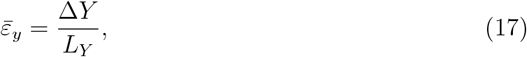

where *L*_*Y*_ is the height of the structure. The normalized displacement *S*(*X, Y*) is then scaled to physical displacement as show in Equation 18:

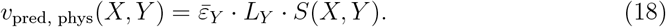

From the reconstructed displacement field, the strain components are computed using Equations (19–21):

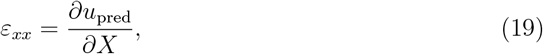

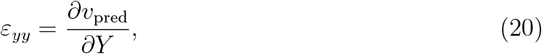

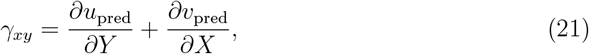

where *ε*_*xx*_ and *ε*_*yy*_ are normal strains along the *X* and *Y* directions, and *γ*_*XY*_ is the engineering shear strain. *X* and *Y* denote the spatial Cartesian coordinates, and *∂/∂X* and *∂/∂Y* represent the corresponding partial derivatives.

Assuming linear elastic material behavior under plane stress conditions, the corresponding stress components (*σ*_*xx*_, *σ*_*yy*_, and *τ*_*xy*_) are computed using Hooke’s law, as expressed in Equations (22–24):

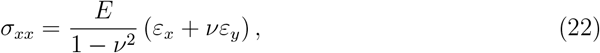

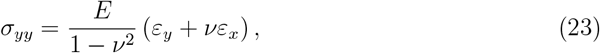

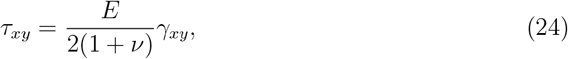

where *σ*_*xx*_ and *σ*_*yy*_ denote the normal stress components acting along the *X* and *Y* directions, respectively, while *τ*_*xy*_ represents the in-plane shear stress component. *E* is Young’s modulus, and *ν* is Poisson’s ratio.

In nonlinear materials, constitutive models are applied in post-processing to transform the predicted displacement or strain distribution into realistic stress fields as in Equation 25:

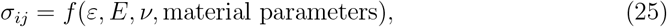

where *f* represents the nonlinear material relationship, which include material parameters such as yield stress, where *σ*_*ij*_ is the in-plane stress tensor with components *σ*_*xx*_, *σ*_*yy*_, and *τ*_*xy*_, representing the normal stresses along the *X*- and *Y* -directions and the in-plane shear stress, respectively. The strain tensor *ε* includes the normal strains *ε*_*xx*_, *ε*_*yy*_ and the engineering shear strain *γ*_*xy*_. The function *f* defines the nonlinear relationship between stress and strain, incorporating material parameters such as yield stress, and the tangent (strain hardening) modulus. In two dimensions, all stress components are coupled through the nonlinear constitutive law, meaning that a change in any strain component can influence multiple stress components. This formulation post-processes AI-BioMech–predicted displacement or strain fields to compute realistic 2D stress distributions. These distributions capture nonlinear material effects, including bilinear isotropic hardening and plastic deformation.

By integrating the predicted color maps with material properties, the network efficiently captures complex spatial patterns without being constrained by the governing physics during training, while still producing quantitatively meaningful stress and strain distributions, instead to enforce physical laws during training, this approach provides computational efficiency and greater flexibility in handling multiple materials and highly irregular cellular geometries.

### 2.5 Evaluation methods

To validate the predicted color maps generated by our semantic segmentation model, we employed several standard evaluation metrics that quantify the accuracy and effectiveness of pixel-wise predictions [55]. These metrics assess not only overall correctness but also class-specific performance, providing a comprehensive measure of how well the model captures the stress and displacement patterns.

**Pixel Accuracy (PA)** calculates the percentage of correctly classified pixels across the entire image, as expressed by Equation 26:

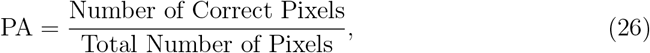

which provides a straightforward measure of the model’s overall correctness.

**Mean Pixel Accuracy (mPA)** measures the average pixel accuracy across all classes, as described by Equation 27:

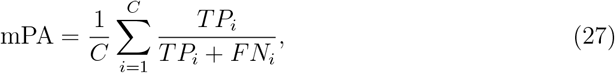

where *C* is the number of classes, *TP*_*i*_ is the number of true positives for class *i*, and *FN*_*i*_ is the number of false negatives for class *i*. This metric is particularly useful for evaluating model performance in the presence of class imbalance, ensuring that smaller or less frequent classes are accurately predicted.

To measure the overlap between the predicted and true segmentation masks, we used the **Mean Intersection over Union (mIoU)**, as defined in Equation 28:

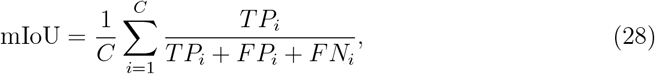

where *FP*_*i*_ represents the false positives for class *i*. The mIoU provides a comprehensive view of the model’s ability to segment each class accurately, accounting for both over- and under-segmentation.

Finally, the **Dice Similarity Coefficient (DSC)** measures the similarity between the predicted and ground truth masks by calculating twice the intersection of the sets divided by the sum of their sizes is described by Equation 29:

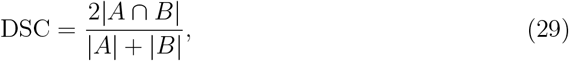

where *A* and *B* represent the predicted and true segmentation masks, respectively. The DSC is particularly sensitive to small structures and helps evaluate how accurately fine stress regions are captured. Together, these evaluation metrics serve to rigorously validate the predicted color maps, ensuring that the DCNN model accurately identifies and segments stress and displacement regions in cellular structures.

#### 3.5.1 Experimental validation of predicted stress–strain behavior

Once the model demonstrated satisfactory performance in both validation and generalization tests, 2D extruded cellular structures were fabricated for experimental validation. Polylactic Acid (PLA) samples inspired by wood and sponge microarchitectures were produced, with ten replicates of each structure type. The PLA filament used was 1.75 mm in diameter with a recommended printing temperature of 195–235^°^C and a bed temperature range of 55–75^°^C. The samples were printed using a Bambu Lab X1 Mini 3D printer employing fused deposition modeling (FDM) technology. A 0.4 mm nozzle was used with a layer height of 0.2 mm, 100% infill, and a print speed of 50 mm/s. The printing orientation was set such that the extrusion direction aligned with the *Z*-axis of the print bed. The bed temperature was maintained at 60^°^C and the nozzle temperature at 210^°^C throughout the printing process.No post-processing or heat treatment was performed on the printed parts. To approximate the simulated boundary conditions, compressive loading was applied from the top edge, corresponding to plane stress conditions in the structure. Compression tests were conducted using an Instron 3369 under displacement-controlled loading, using a 50kN load cell, applying a total displacement equal to 20% of the specimen height at a loading rate of 0.05 mm/sec. Full-field displacement and strain distributions were captured using Digital Image Correlation (DIC), and experimental stress–strain curves were derived from these measurements. These experimental results were then compared with numerical predictions obtained from the DCNN-generated displacement maps, providing quantitative validation of both the segmentation results and the post-processed stress predictions.

Overall, this methodology establishes an integrated pipeline for AI-BioMech framework validation, combining image-based FEA, deep learning–based segmentation, and experimental testing. By linking the DCNN-predicted displacement fields with measured stress–strain responses, the framework ensures that the modeled mechanical behavior is both physically meaningful and experimentally verifiable.

## 3 Results and Discussion

### 3.1 Benchmarking Outcomes of the Synthetic Dataset

In our study, we used the pretrained ResNet-101 network to extract visual features from each image. Each image generates a 2048-dimensional feature vector, and for the set of 200 reference images, this results in a total of 200 × 2048 feature vectors. Each test image was then compared against all 200 reference feature vectors to compute both cosine similarity and Euclidean distance, providing quantitative measures of structural and geometric similarity. Across the entire synthetic dataset of 350,000 images, this process produced a total of 70,000,000 similarity values, which were subsequently summarized and visualized using box-and-whisker plots to evaluate the distribution and range of feature alignment with the reference images.

Figure 7 provides a qualitative comparison of feature representations between real and synthetic datasets across different cellular structures. The cross-domain feature correlation maps showcase how effectively the synthetic dataset captures the structural cues present in real samples, with the color scale indicating the degree of similarity: red regions represent strong feature alignment, whereas blue regions highlight areas of lower correspondence. These low-similarity regions suggest the presence of additional feature variations in the synthetic data that are not commonly observed in natural cellular structures such as wood, sponge, and bone. Rather than being a limitation, this variability can be beneficial for model training, as it introduces feature diversity that encourages better generalization and reduces overfitting to specific structural patterns [58].

**Figure 7.**
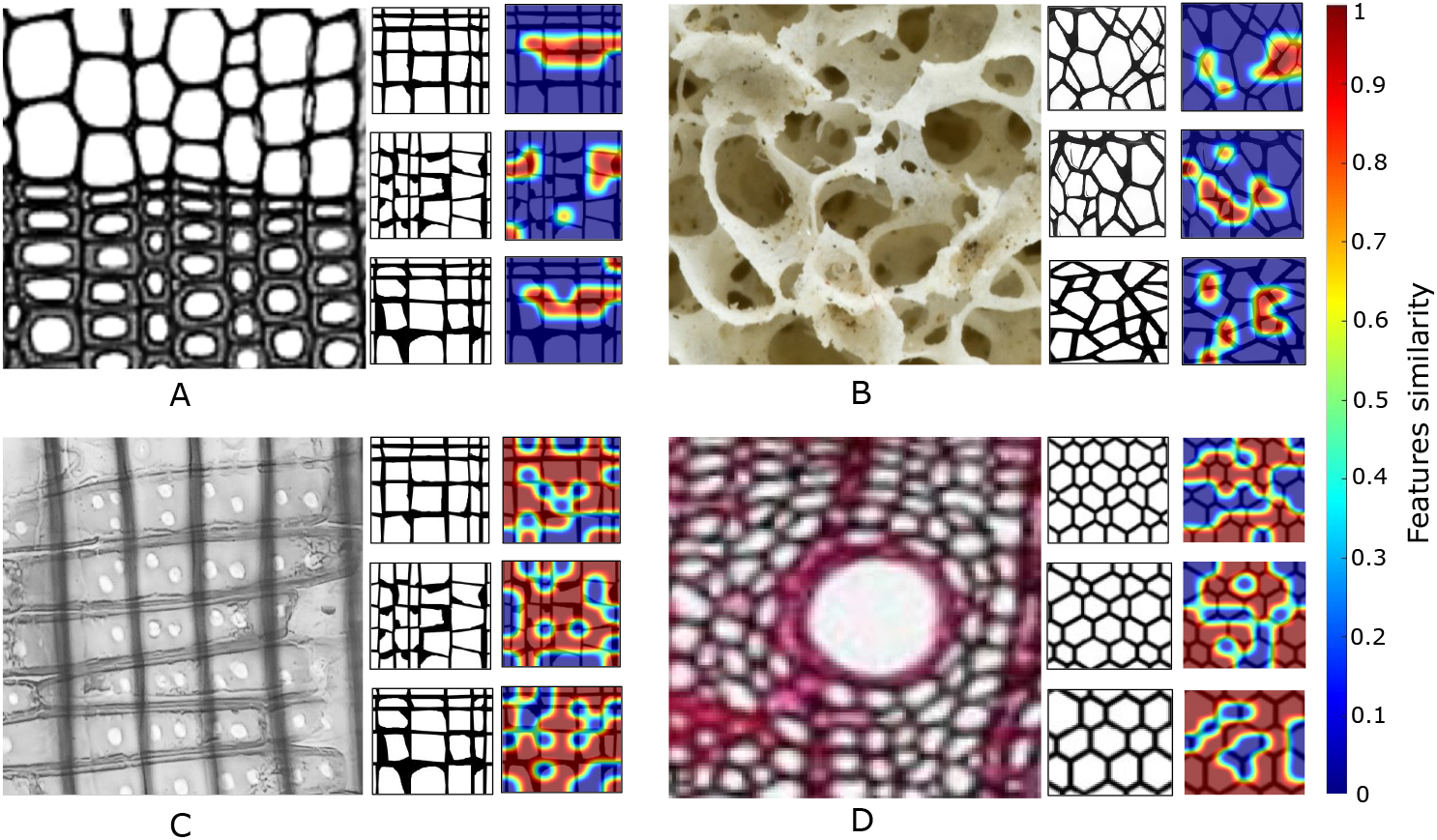
Map visualization showing the features correlation between real and synthetic dataset samples. (A) illustrates Gymnosperm stem soft wood in *Pinus* cellular structure with a cosine similarity of 0.4 to the synthetic data, while (B) shows a bone cellular structure with a cosine similarity of 0.5. (C) depicts Calabrian pine (*Pinus brutia*) wood structure with a higher similarity of 0.8, and (D) shows a Quercus stem wood cellular structure with a similarity of 0.7. These maps highlight varying levels of alignment between real and synthetic samples, with higher cosine similarity indicating stronger correlation. Original microstructure images (used to generate real samples) were obtained under CC0 /CC BY licenses [56] [57].

As shown in Figure 8, synthetic and real-world data samples were compared using cosine similarity and Euclidean distance to evaluate how closely the synthetic dataset reflects real-world features.The cosine similarity dataset represented predominantly strong correlation, with approximately 60% of synthetics dataset samples falling within a high-similarity range (0.70–1.00), 25% in a moderate similarity region (0.41–0.69), and the remaining 15% representing low similarity (0–00.40). This distribution emphasizes cases where synthetic dataset fields closely match the ground truth, while still incorporating moderate and weak alignment scenarios.

**Figure 8.**
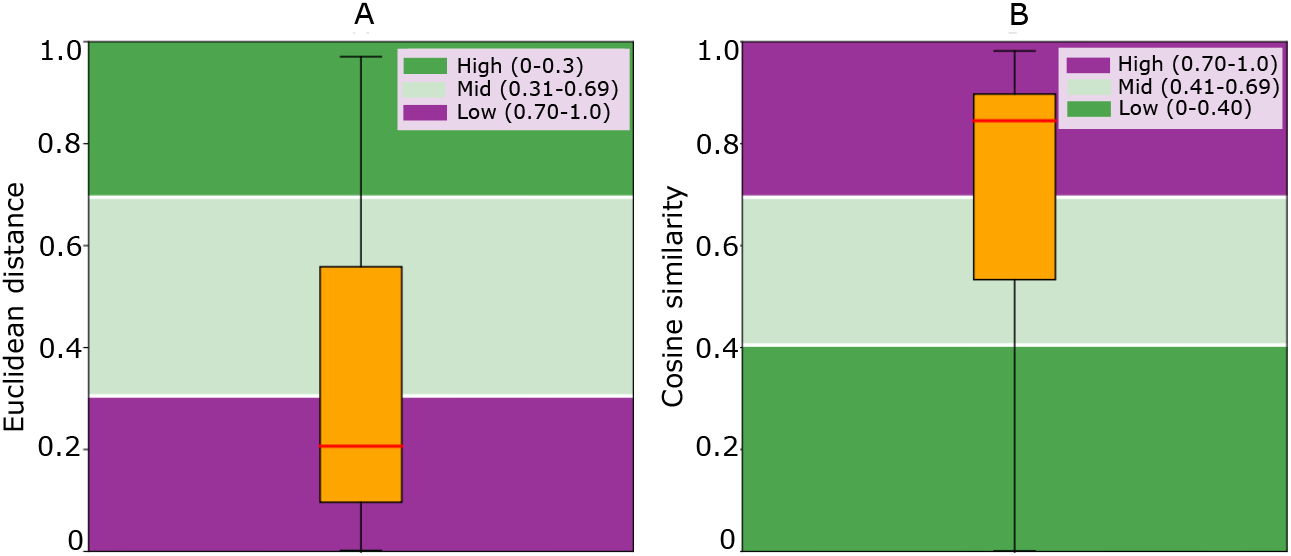
Illustration of the Euclidean distance and cosine similarity distributions between synthetic and real-world samples, demonstrating how the synthetic dataset aligns with real-world data.The real-world dataset contains 200 samples, while the synthetic dataset includes 350,000 samples, resulting in a total of 70,000,000 pairwise similarity values used for analysis.

Conversely, the Euclidean distance metric serves as an inverse measure of similarity, where lower values indicate a higher degree of correlation between synthetic and real world features. In this dataset, approximately 70% of synthetics samples fall within the low-distance range (0–0.31), representing highly correlated with minimal spatial deviations. Around 20% of samples lie in the moderate-distance range (0.31–0.69), indicating intermediate correlation, while 10% of samples occupy the high-distance range (0.70–1.00), corresponding to poorly correlated with larger discrepancies from the reference. This distribution mirrors real-world variability, showing that most synthetic samples closely match the real world data, while still capturing occasional deviations.

Based on the feature comparison analysis, approximately 85% of the synthetic dataset exhibits high and moderate correlation with real-world samples, as indicated by cosine similarity and Euclidean distance metrics. This strong alignment suggests that the synthetic data effectively captures the key structural characteristics of real cellular materials, as projected here from both wood and sponge architectures. Consequently, the dataset provides a representative and diverse set of features that can be reliably used for training and validating machine learning models. The remaining 15% of samples with lower correlation still contribute variability that reflects real-world noise and structural diversity, further enhancing the robustness of model training. Therefore, the results indicate that the synthetic dataset is sufficiently accurate and comprehensive to serve as a practical surrogate for real-world data in predictive tasks.

### 3.2 Performance of optimized transfer learning models

Following validation of the synthetic dataset, model training was performed on a workstation equipped with a single NVIDIA GPU, using MATLAB for data preprocessing and network implementation. The datasets were processed to conform to the input specifications of the pre-trained backbone networks. Three pre-trained architecture: ResNet50, ResNet101, and Inception-ResNetV2, were employed as feature extractors within the proposed encoder-decoder framework to perform semantic segmentation of von Mises stress, shear stress, and X/Y displacement distributions in cellular structures. The dataset, comprising 350,000 images, was split into 70% for training, 15% for testing, and 15% for validation to ensure proper evaluation of model performance. Various hyperparameter settings, including batch size, learning rate, and optimizer type, were evaluated to optimize network training. Figure 9 illustrates the effect of these variations on model performance.

**Figure 9.**
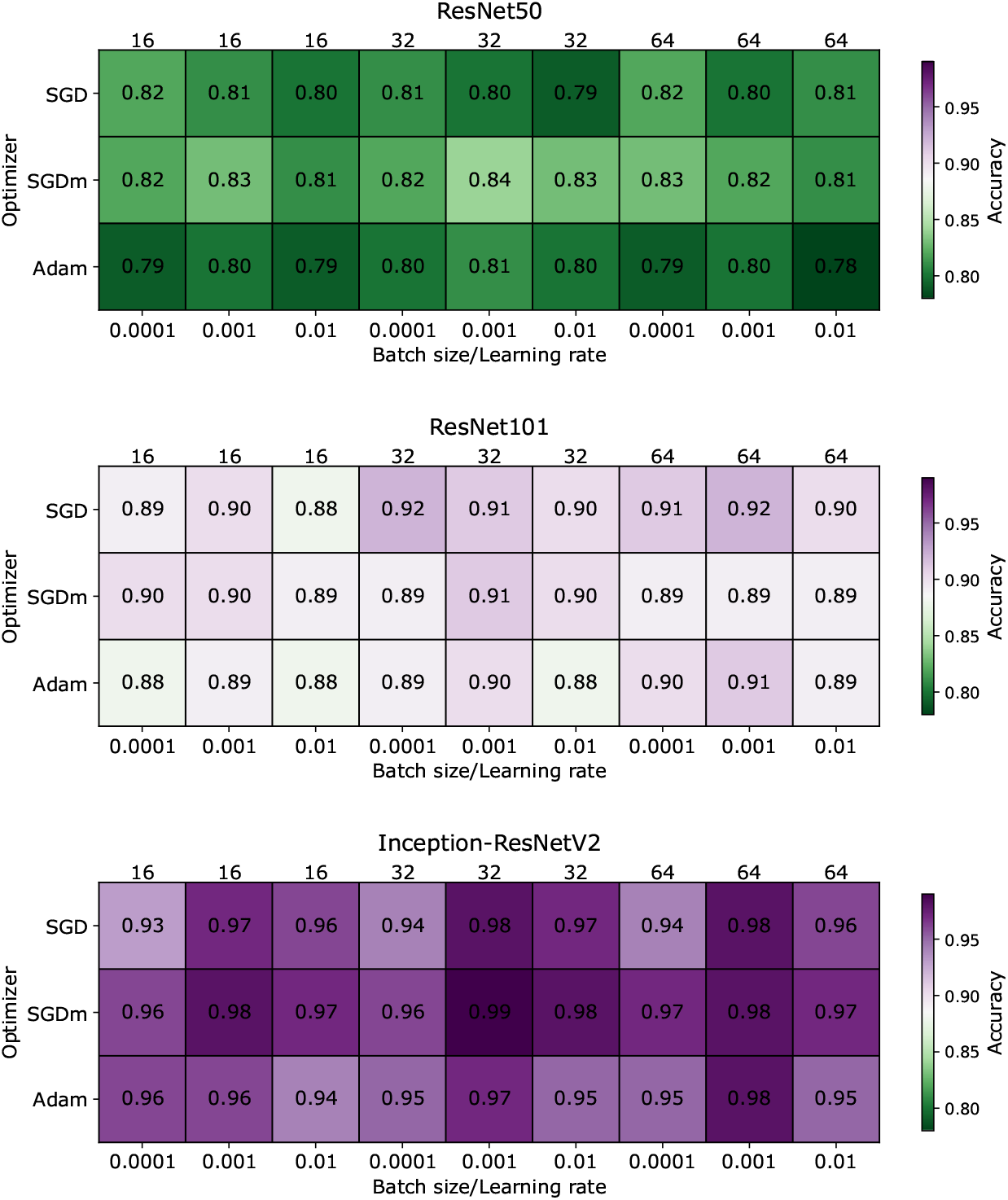
Comparison of model accuracy for ResNet50, ResNet101, and Inception- ResNetV2 across different hyperparameter settings, including learning rate, batch size, and optimizer

Among the backbone architectures tested, ResNet50 exhibited moderate performance, with accuracies ranging from 0.79 to 0.84. The model achieved its highest validation accuracy of 0.84 using a learning rate of 0.001, a batch size of 32, and the SGD optimizer, indicating that mid-range learning rates and moderate batch sizes are optimal for this architecture. Training progress and performance metrics are presented in Appendix A, Figure A1, which shows the ResNet50 validation accuracy reaching 84% and a corresponding loss of 0.4% over 100 epochs.

Building on this, ResNet101 demonstrated improved performance, achieving accuracies between 0.88 and 0.91. The highest validation accuracy of 0.91 was obtained using the SGDm optimizer, a learning rate of 0.001, and a batch size of 32, highlighting the advantage of a more complex architecture and adaptive optimization for enhanced feature extraction. Appendix A, Figure A2, shows the ResNet101 training graphs with an optimal validation accuracy of 91% and loss of 0.2% over 100 epochs. The Inception-ResNetV2 architecture achieved the highest overall accuracy, ranging from 0.93 to 0.99. The peak accuracy of 0.99 was obtained using the SGDm optimizer, a learning rate of 0.001, and a batch size of 32, demonstrating that this architecture can most effectively leverage the synthetic dataset to predict von Mises stress, shear stress, and displacement fields when combined with appropriate hyperparameters. Appendix A, Figure A3, provides the Inception-ResNetV2 training graphs, depicting an optimal validation accuracy of 99.98% and a loss of 0.0005% over 100 epochs.

The validation loss trends across the three architectures reflect their relative effectiveness in learning the target stress and displacement features. ResNet50 exhibited the highest validation loss of 0.40%, indicating moderate prediction accuracy and residual error in capturing complex stress and displacement patterns. ResNet101 demonstrated improved performance, with a reduced validation loss of 0.20%, reflecting enhanced generalization and more accurate feature extraction due to its deeper architecture. Inception-ResNetV2 achieved the lowest validation loss of 0.005%, indicating highly accurate predictions. Its superior performance can be attributed to the combination of increased depth and the inception modules, which enable the network to simultaneously capture fine-grained local features and broader global patterns more effectively than the other models.

Comparing the training times of the three architectures highlights the trade-off between predictive accuracy and computational cost (see Figure 10). ResNet50 achieved reasonable accuracy while requiring the shortest training time, ranging from 12.05 to 15.24 hours. ResNet101 required longer training, between 14.20 and 17.30 hours, reflecting the increased computational demand of its deeper architecture. Inception-ResNetV2 provided the highest predictive accuracy, but this came at the expense of the longest training times, ranging from 17.10 to 21.21 hours. These results indicate that while deeper and more complex architectures improve model performance, they also demand greater computational resources, emphasizing the need to balance accuracy and efficiency when selecting an appropriate network for large-scale stress and displacement predictions.

**Figure 10.**
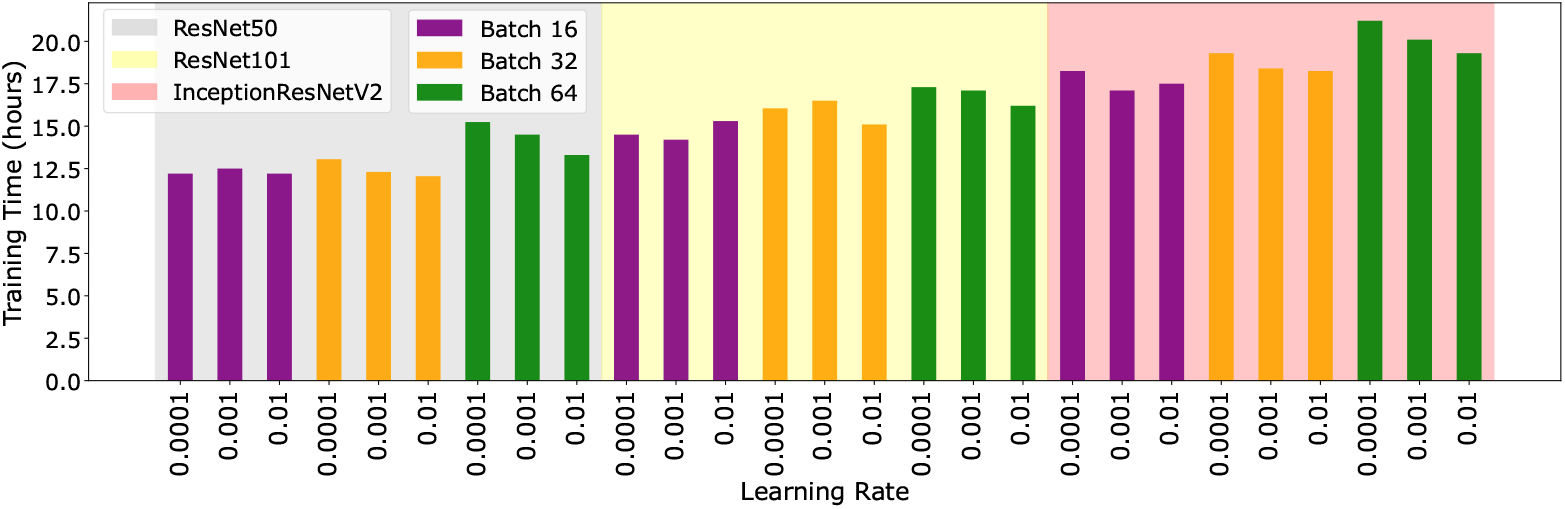
Training times for ResNet50, ResNet101, and Inception-ResNetV2 across different learning rates and batch sizes. The figure illustrates that while Inception-ResNetV2 achieves the highest accuracy, it requires longer training compared to the other models, highlighting the trade-off between performance and computational cost

### 3.3 Segmentation performance evaluation

To assess the segmentation performance of the proposed encoder–decoder architecture with three fine-tuned architectures: ResNet50, ResNet101, and Inception-ResNetV2, both quantitative and qualitative evaluations were carried out. The qualitative analysis was conducted using predicted outputs X-displacement, Y-displacement, von Mises stress, and shear stress colormaps as shown in Figure 11, 12,13, and 14 respectively. Across these visual results, the Inception-ResNetV2 backbone consistently delivered the most accurate and continuous stress field representations, characterized by sharper boundaries and reduced noise. ResNet101 also produced visually acceptable segmentation, although slight blurring was observed in areas containing steep stress gradients. In comparison, ResNet50 displayed the weakest performance, with more coarse predictions and less distinct stress transitions, especially in regions with high stress and displacement variations.

**Figure 11.**
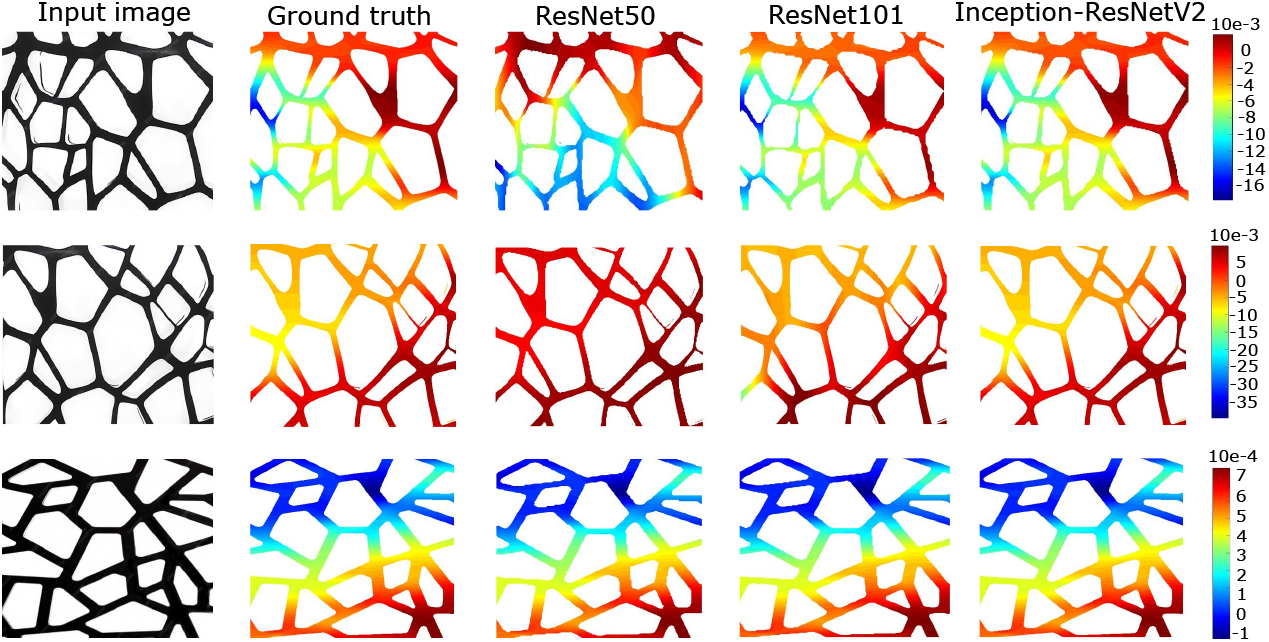
Comparison of X-displacement predictions for a cellular structure using fine-tuned deep learning architectures: ResNet50, ResNet101, and Inception-ResNetV2, alongside the corresponding ground truth. A vertical displacement of 0.001 mm was applied at the top boundary, while the bottom edge was fixed, with Young’s modulus of 210 GPa and Poisson’s ratio of 0.3. These reference conditions provide the basis for evaluating semantic segmentation–based X-displacement predictions from the three models

**Figure 12.**
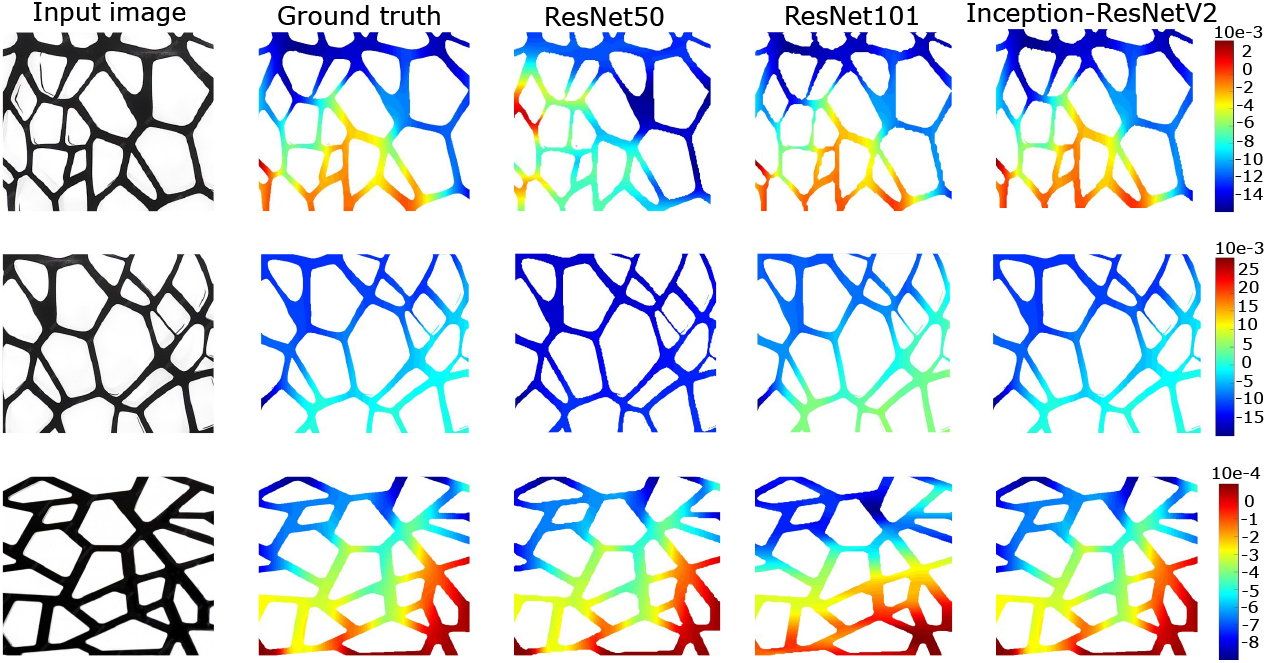
Comparison of Y-displacement predictions for a cellular structure using fine-tuned deep learning architectures: ResNet50, ResNet101, and Inception-ResNetV2, alongside the corresponding ground truth. A vertical displacement of 0.001 mm was applied at the top boundary, while the bottom edge was fixed, with Young’s modulus of 210 GPa and Poisson’s ratio of 0.3. These reference conditions provide the basis for evaluating semantic segmentation–based Y-displacement predictions from the three models

**Figure 13.**
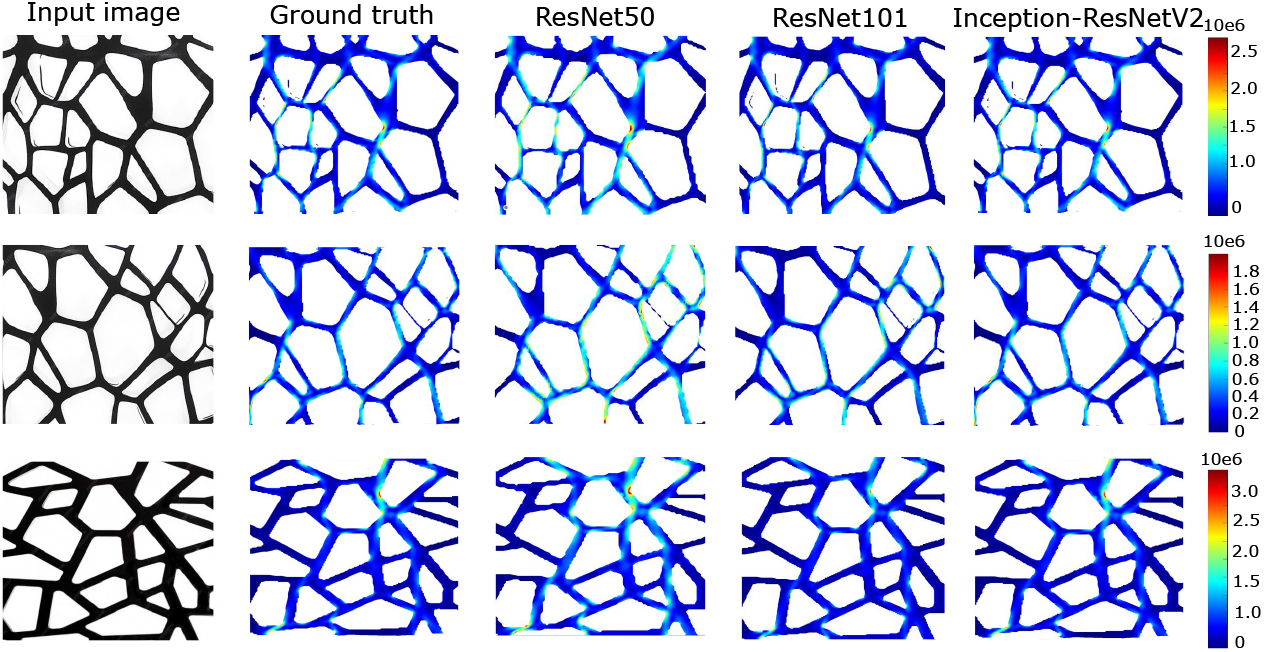
Comparison of von Mises stress predictions for a cellular structure using fine-tuned deep learning architectures: ResNet50, ResNet101, and Inception-ResNetV2, alongside the corresponding ground truth. A vertical displacement of 0.001 mm was applied at the top boundary, while the bottom edge was fixed, with Young’s modulus of 210 GPa and Poisson’s ratio of 0.3. These reference conditions provide the basis for evaluating semantic segmentation–based von Mises stress predictions from the three models

**Figure 14.**
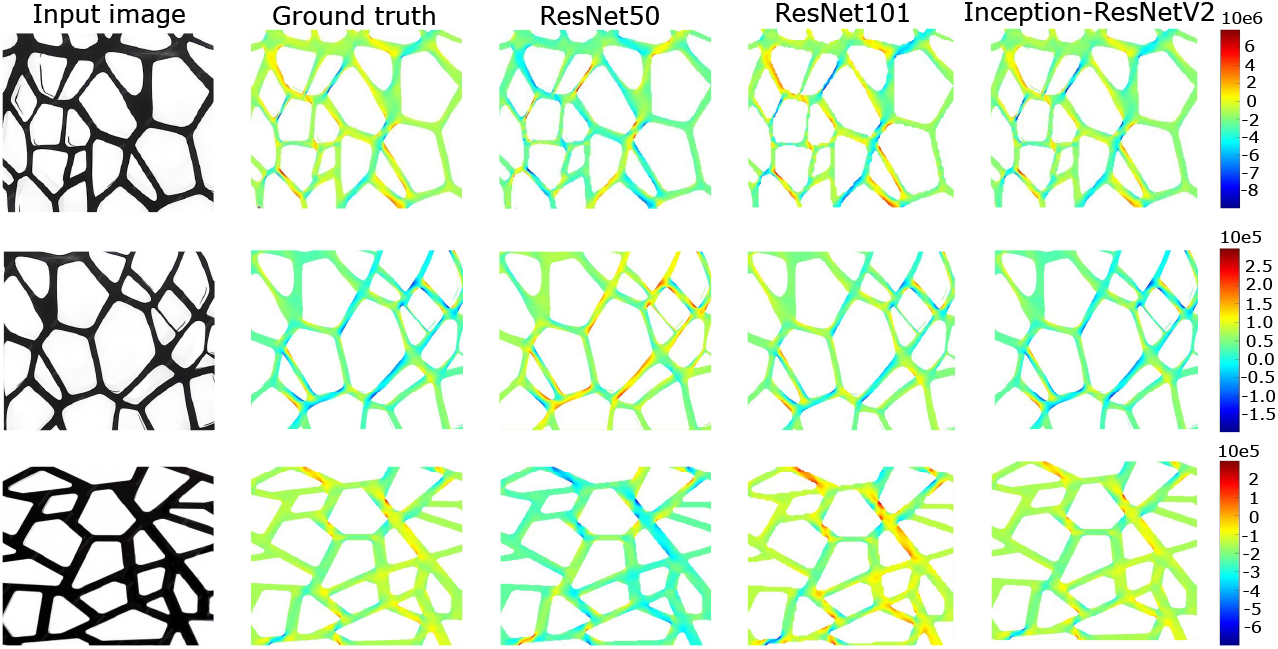
Comparison of shear stress predictions for a cellular structure using fine-tuned deep learning architectures: ResNet50, ResNet101, and Inception-ResNetV2, alongside the corresponding ground truth. A vertical displacement of 0.001 mm was applied at the top boundary, while the bottom edge was fixed, with Young’s modulus of 210 GPa and Poisson’s ratio of 0.3. These reference conditions provide the basis for evaluating semantic segmentation–based shear stress predictions from the three models

Table 1 details the quantitative performance comparison evaluation of three fine-tuned architectures, using four segmentation performance metrics: Pixel Accuracy (PA), mean Pixel Accuracy (mPA), mean Intersection-over-Union (mIoU), and Dice Similarity Coefficient (DSC). These metrics collectively measure pixel-wise correctness, class-wise accuracy, overlap between predicted and ground-truth masks, and similarity in segmented regions. The results clearly show that Inception-ResNetV2 consistently outperforms both ResNet50 and ResNet101 across all evaluation indicators. It achieves the highest PA (99.05%), mPA (98.65%), mIoU (98.40%), and DSC (98.10%) for stress and X/Y displacement prediction. This outstanding performance suggests that the hybrid design of Inception modules with residual connections enables richer multi-scale feature extraction and more robust convergence behavior, leading to highly precise segmentation outputs.

**Table 1.**
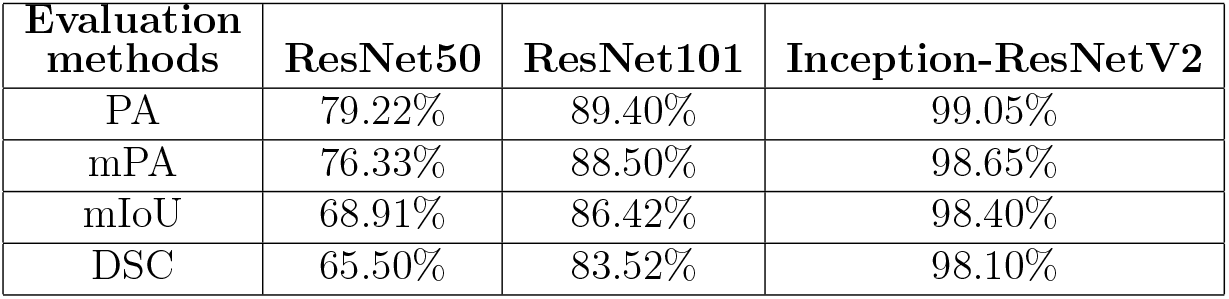
Quantitative evaluation of segmentation performance of ResNet50, ResNet101, and Inception-ResNetV2 using PA, mPA, mIoU, and DSC.

| **Evaluation methods** | **ResNet50** | **ResNet101** | **Inception-ResNetV2** |
| --- | --- | --- | --- |
| PA | 79.22% | 89.40% | 99.05% |
| mPA | 76.33% | 88.50% | 98.65% |
| mIoU | 68.91% | 86.42% | 98.40% |
| DSC | 65.50% | 83.52% | 98.10% |

Following the selection of Inception-ResNetV2 as the best-performing model, its generalization capability was further validated using 500 previously unseen test samples. This evaluation was conducted to measure the model’s robustness and reliability when applied to real-world data beyond the training set.

As presented in Table 2, the model maintained consistently strong performance across 90%, 95%, and 99% confidence levels, reflecting stable and dependable behavior under different statistical conditions. At the 90% confidence interval, PA ranged from 97.20–99.05%, mPA from 97.31–98.22%, mIoU from 96.20–98.43%, and DSC from 97.20–98.55%. Similar performance was observed at the 95% confidence level, with PA between 96.22–98.99%, mPA 96.40–98.32%, mIoU 95.32–98.34%, and DSC 96.40–98.07%. Even under the stricter 99% confidence bounds, all metrics remained above 94%, with PA between 96.22–99.01%, mPA 94.21–98.10%, mIoU 94.23–98.10%, and DSC 94.32–98.04%. Although the ranges widen slightly at higher confidence levels due to increased statistical uncertainty, the narrow intervals overall indicate excellent generalization, low performance variance, and high segmentation precision. These findings confirm Inception-ResNetV2 as a robust, stable, and highly effective backbone for practical real-world deployment.

**Table 2.** Performance evaluation of the Inception-ResNetV2 model, selected as the optimal backbone, showing metrics at 90%, 95%, and 99% confidence levels based on 500 real word test samples.

| **Evaluation Methods** | **Confidence levels** |  |  |
| --- | --- | --- | --- |
|  | **90%** | **95%** | **99%** |
| PA | (97.20-99.05)% | (96.22-98.99)% | (96.22-99.01)% |
| mPA | (97.31-98.22)% | (96.40-98.32)% | (94.21-98.10)% |
| mIoU | (96.20-98.43)% | (95.32-98.34)% | (94.23-98.10)% |
| DSC | (97.20-98.55)% | (96.40-98.07)% | (94.32-98.04)% |

### 3.4 Physics-informed post-processing of model predictions

The mechanical response of the wood material was predicted by integrating the governing physics equations, with the resulting von Mises stress distribution visualized as a color map (Figure 15). Each color in the map corresponds to a local stress value, which was then converted into strain by incorporating the material properties of wood, specifically a Young’s modulus of *E* = 10 GPa and a Poisson’s ratio of *ν* = 0.3. A prescribed displacement of 0.4% of the total length of the cellular structure was applied along the top edge to induce stress and strain in the material.

**Figure 15.**
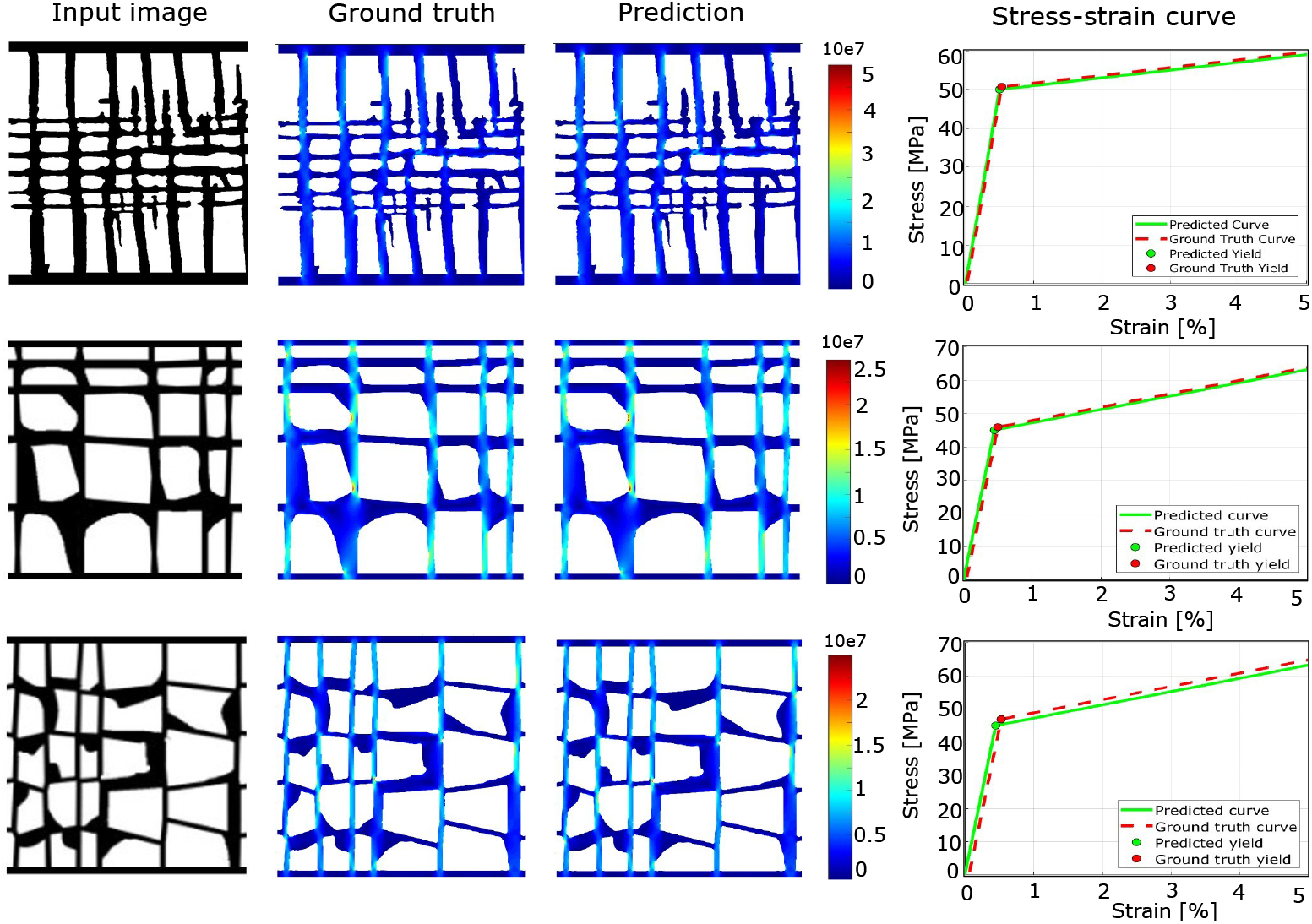
Predicted von Mises stress distribution of the wood material obtained by integrating the governing physics equations with the model predictions. A prescribed displacement of 0.4% of the total length was applied along the top edge. Material properties used include a Young’s modulus of *E* = 10 GPa and a Poisson’s ratio of *ν* = 0.3. The bilinear constitutive model captures both elastic and plastic responses, with yielding occurring at 50 × 10^6^ Pa. Color contours indicate the local stress/strain distribution across the irregular cellular structure

By mapping the stress values from the color gradient to the material’s constitutive behavior, the local stress-strain response was reconstructed using a bilinear constitutive model, capturing both elastic and plastic regions. Yielding was defined at a stress threshold of 50 MPa. The predicted stress-strain behavior closely matches the ground truth measurements: for the top sample, the ground truth yield strain was 0.49% and the predicted value was 0.51%; for the middle sample, the values were 0.45% (ground truth) and 0.48% (predicted); and for the bottom sample, 0.48% and 0.49%, respectively. These results indicate that the model accurately captures the onset of plastic deformation across different locations in the structure.

These results demonstrate that integrating material properties with the color-mapped stress field allows for accurate prediction of both elastic and plastic responses. The color map also provides a visual representation of localized stress and strain distributions across the irregular cellular structure of wood, highlighting regions approaching or exceeding the yield threshold. Minor deviations between predicted and ground truth values can be attributed to simplifications in the constitutive model and the inherent variability of the wood microstructure.

### 3.5 Experimental validation

In this study, 3D-printed PLA samples with wood and sponge cellular structures were tested to validate our physics-based integration against experimental results and to compare the outcomes of our semantic segmentation framework with Digital Image Correlation (DIC) measurements. The experiments aimed to characterize the stress–strain behavior of the cellular structures under compression, with particular focus on both the elastic and plastic regions. Specimens were subjected to uniaxial compression under a plane strain condition, consistent with our prediction model, which analyzes 2D images under the assumption of planar stress–strain behavior.

Experiments were conducted on an Instron 3369 universal testing machine using a 50 kN load cell, as shown in Figure 16. The specimen dimensions were uniform at 5.5 cm × 5.5 cm × 3.5 cm (length × height × width). For statistical reliability, ten specimens of each structure (wood and sponge) were tested. The resulting stress–strain curves show the average response, and the shaded region represents the spread, highlighting the variability used to analyze deviations between specimens.

**Figure 16.**
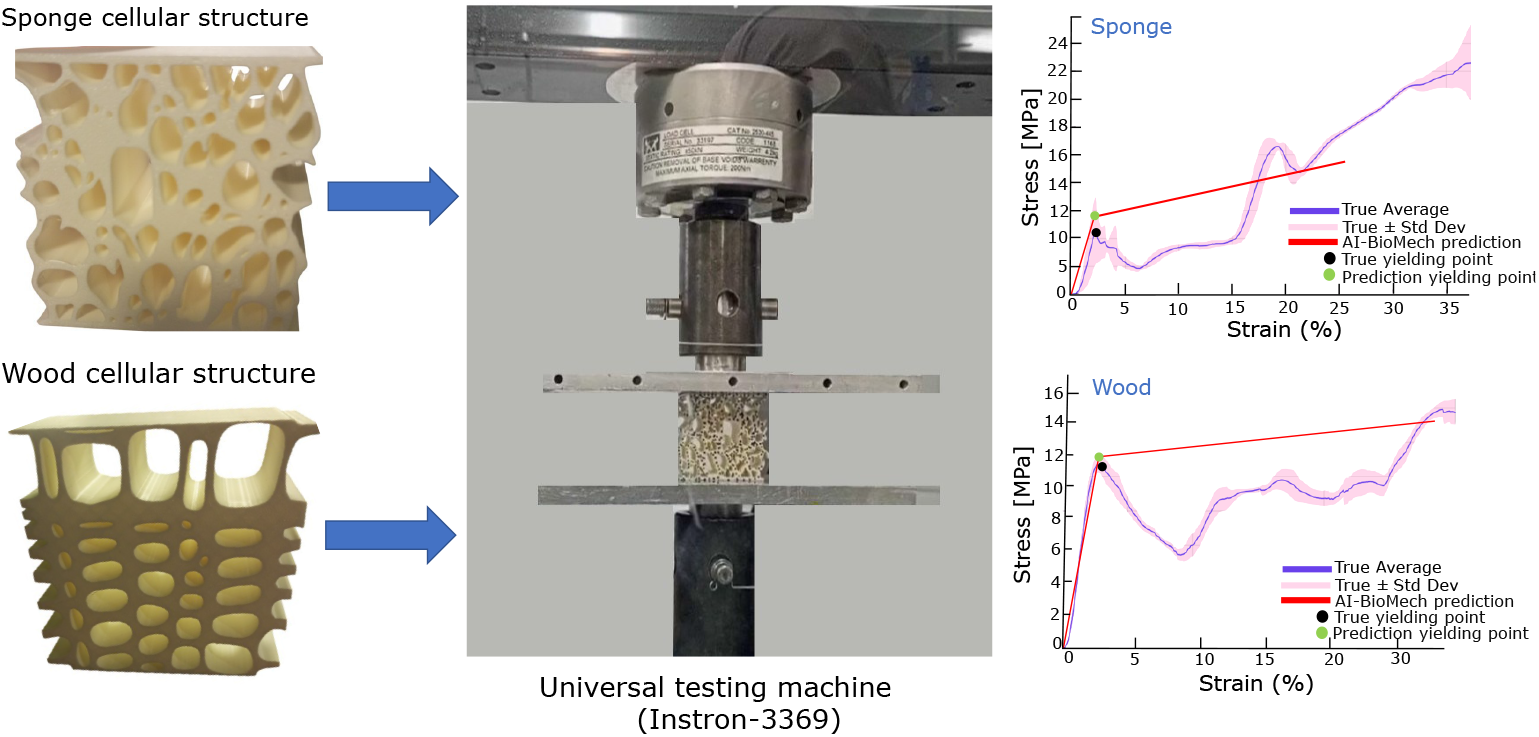
3D PLA samples with wood and sponge cellular structures were tested to validate our physics-based integration against experimental results and to compare semantic segmentation outcomes with Digital Image Correlation (DIC) measurements. Ten specimens samples were tested for each structure (total=20 samples) with dimensions of 5.5 cm × 5.5 cm × 3.5 cm (length × height × width), and the plain stress-strain response was calculated from the measured load and extension data. Tests were performed using an Instron 3369 universal testing machine under a 50kN load cell. Each curve represents the mean response of ten specimens, with the shaded region indicating the spread of the data as standard deviation

As shown in Figure 16, the experimental stress–strain curves exhibit continuous, non-linear behavior beyond the yield point. In contrast, our model produces an approximately bilinear response, derived from the color map gradient of the integrated governing equations. The model is based on a Young’s modulus of 2.5 × 10^9^ Pa and a yield strength of 60 × 10^6^ Pa. The adopted material parameters are consistent with literature values for PLA, which typically report a Young’s modulus in the range of 2–3.5 GPa and a tensile yield strength between 50–70 MPa. We compared the predicted and experimental curves mainly to assess how well the model captures the yielding point. The sponge cellular structure, the experimentally measured yield stress is 10.5 MPa at a strain of 2.3%, while the model predicts a yield stress of 12 MPa at 2.1% strain. For the wood cellular structure, the experimental yield stress is 11 MPa at 2.2% strain, whereas the model predicts 11.50 MPa at 2% strain. In both cases, the predicted results lie within the spread of the experimental measurements, demonstrating that the bilinear model captures the experimental yielding behavior accurately.

In the elastic region, our bilinear model closely replicates the initial linear behavior of the experimental stress–strain curves, accurately capturing both the slope and the approximate yielding point. In the plastic region, although the experimental response exhibits nonlinearity, the model effectively follows the overall trend, with the tangent slope of the predicted curve mirroring the experimental behavior. Furthermore, the predicted stresses remain within the range of experimental variability, demonstrating that the prediction model provides a reasonable approximation of the post-yield response.

### 3.6 Digital Image Correlation Analysis

The displacement and strain fields of the 3D cellular specimens, including wood and sponge structures, were measured using 2D Digital Image Correlation (DIC) to validate the predictions of our model and the outcomes of the semantic segmentation framework. The analysis was conducted with the **Ncorr toolbox** in MATLAB, which computes displacement and strain fields from high-resolution images captured before and after deformation. To ensure accurate measurements, binary masks were created to define regions of interest (ROIs), excluding background and void areas. Within these ROIs, the X- and Y-displacements as well as the strain components *ε*_*xx*_, *ε*_*yy*_, and *ε*_*xy*_ were extracted and combined to calculate the von Mises equivalent stress. This approach enabled the validation of our predicted segmentation results for von Mises stress, shear strain, and displacement by direct comparison with the corresponding DIC-measured fields.

Figure 17 shows the original and deformed shapes of the wood and sponge cellular structures highlighting structural deformation under the applied 50kN load cell, along with the predicted and DIC-measured shear strain (*ε*_*xy*_) colormaps. For the wood structure, high-strain regions are localized around strut junctions and edges, indicating areas of strain concentration, where the prediction model accurately captures the localized high-strain regions around strut junctions and edges, indicating areas of stress concentration. In contrast, the sponge structure exhibits larger overall deformation due to its more compliant open-cell design, with higher strain observed around the thinner cell walls and a more uniform distribution across the structure. A comparison of the displacement and von Mises fields predicted by the CNN model and measured via DIC is provided in Appendix B (Figure A4 and A5).

**Figure 17.**
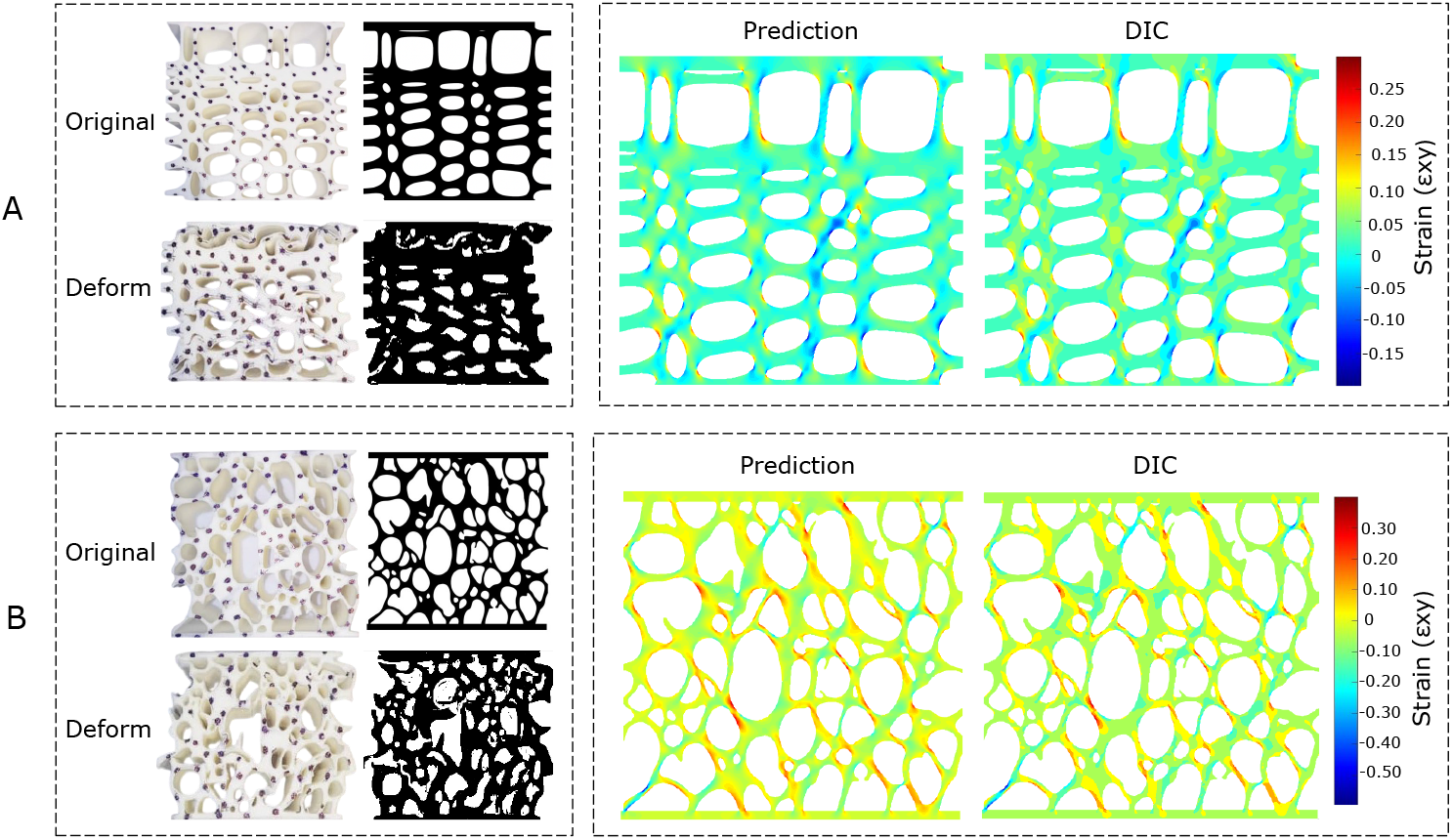
Comparison of original and deformed geometries, and predicted vs experimentally measured shear strain (*ε*_*xy*_) distributions for two 3D-printed cellular structures under 50kN load cell: (A) 3D-printed bone cellular structure and its deformed shape (B) 3D-printed sponge cellular structure and its deformed shape. Predicted fields and DIC-measured colormaps show the shear strain (*ε*_*xy*_) distribution for both cellular structures, illustrating the agreement between CNN predictions and experimental measurements

Overall, the qualitative and quantitative comparison demonstrates that the AI-BioMech framework accurately captures both the displacement and internal strain patterns of the cellular architectures, effectively highlighting how their mechanical response varies with topology.

## 4 Conclusion

In this study, we developed and rigorously validated the AI-BioMech framework, a deep learning–based methodology for predicting displacement, strain, and stress distributions in complex cellular architectures. The framework requires only a single high-resolution 2D image of the structure as input, eliminating the need for extensive experimental measurements or full-scale finite element simulations, thereby significantly reducing computational cost and experimental effort. Large-scale synthetic datasets were used to train deep convolutional neural networks via transfer learning, employing pre-trained backbone architectures including ResNet50, ResNet101, and Inception-ResNetV2. Inception-ResNetV2 emerged as the optimal model, achieving a validation accuracy of 99% due to its deeper architecture and inception modules, which enable simultaneous capture of fine-grained local features and global structural patterns. Hyperparameter optimization further enhanced predictive performance, ensuring robust and reliable results across diverse cellular geometries. Predicted displacement fields were integrated with material constitutive models through a physics-based post-processing step, incorporating both linear and nonlinear stress-strain formulations as appropriate. This approach accurately reconstructed local and global stress-strain distributions, including the onset of plastic deformation, closely matching finite element simulations and experimental measurements. Validation experiments using 3D-printed PLA cellular structures confirmed the framework’s ability to capture both global mechanical response and local strain patterns, demonstrating strong quantitative and qualitative agreement with real-world behavior. The AI-BioMech framework offers several key advantages. It is computationally efficient, able to model materials with different properties, complex geometries, and nonlinear constitutive behaviors. It is a user-friendly software framework, requiring only a single 2D input image. These features make it a versatile tool for materials design, and structural optimization, where rapid, spatially resolved mechanical predictions are critical. Overall, AI-BioMech represents a robust integration of deep learning and physics-based modeling, bridging data-driven predictions with mechanistic validation. By combining high accuracy, efficiency, and accessibility, it provides a reliable platform for the design, evaluation, and optimization of complex cellular structures, enabling researchers and engineers to accelerate innovation in bio-inspired and engineered materials. In the future, the framework could be enhanced to handle 3D material property predictions, facilitating more detailed and realistic modeling of cellular architectures.

## Supporting information

AIBioMech executable

AIBioMech User Manual

## Supplementary Files

- AIBioMech.exe - software executable file
- AIBioMech user manual.pdf - the user manual

## Acknowledgments

The first author (HS) wishes to thank the Higher Education Commission (HEC) Pakistan, for providing the PhD funding that facilitated this research.

## Author Contributions

Conceptualization (PA [lead]); Data curation (HS [lead]); Formal analysis (HS [lead]); Funding acquisition (HS [lead]); Investigation (HS [lead], PA [supporting]); Methodology (HS [lead], PA [supporting], MAD [supporting]); Project administration (PA [lead], MAD [supporting]); Resources (PA [lead]); Software (HS [lead], PA [supporting]); Supervision (PA [lead], MAD [supporting]); Validation (HS [equal], PA [equal]); Visualisation (HS [lead]); Roles/Writing - original draft (HS [lead]); Writing - review and editing (PA [lead], MAD [supporting]).

## A Appendix

### A.1 Training graphs

**Figure A1.**
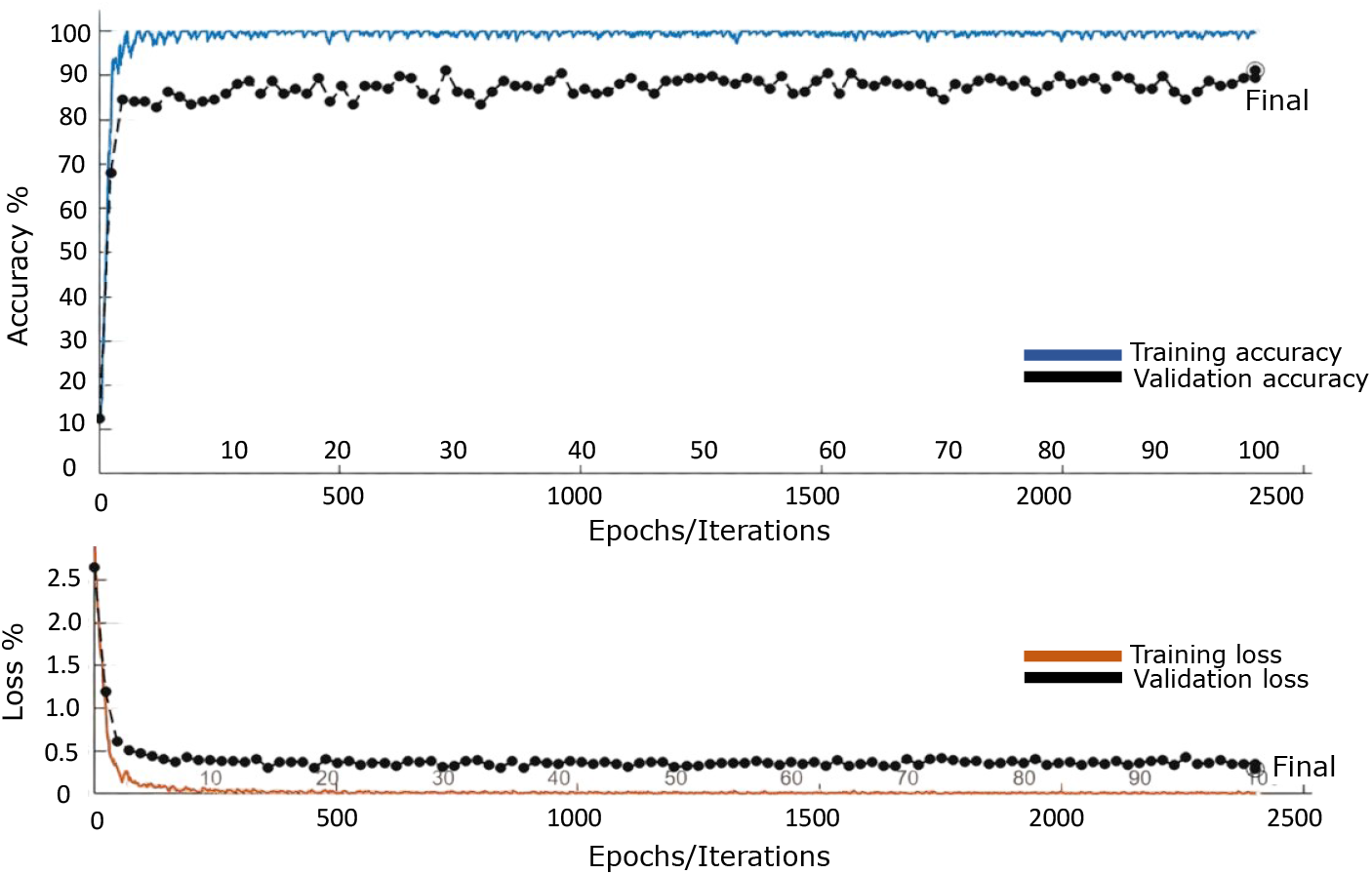
Illustrates training and validation performance of ResNet50. The model achieved an optimal training accuracy of 97% with a training loss of 0.15%, and a validation accuracy of 84% with a validation loss of 0.4%. The model was trained using a learning rate of 0.001, batch size of 32, dropout rate of 0.3, over 100 epochs and 2500 iterations.

**Figure A2.**
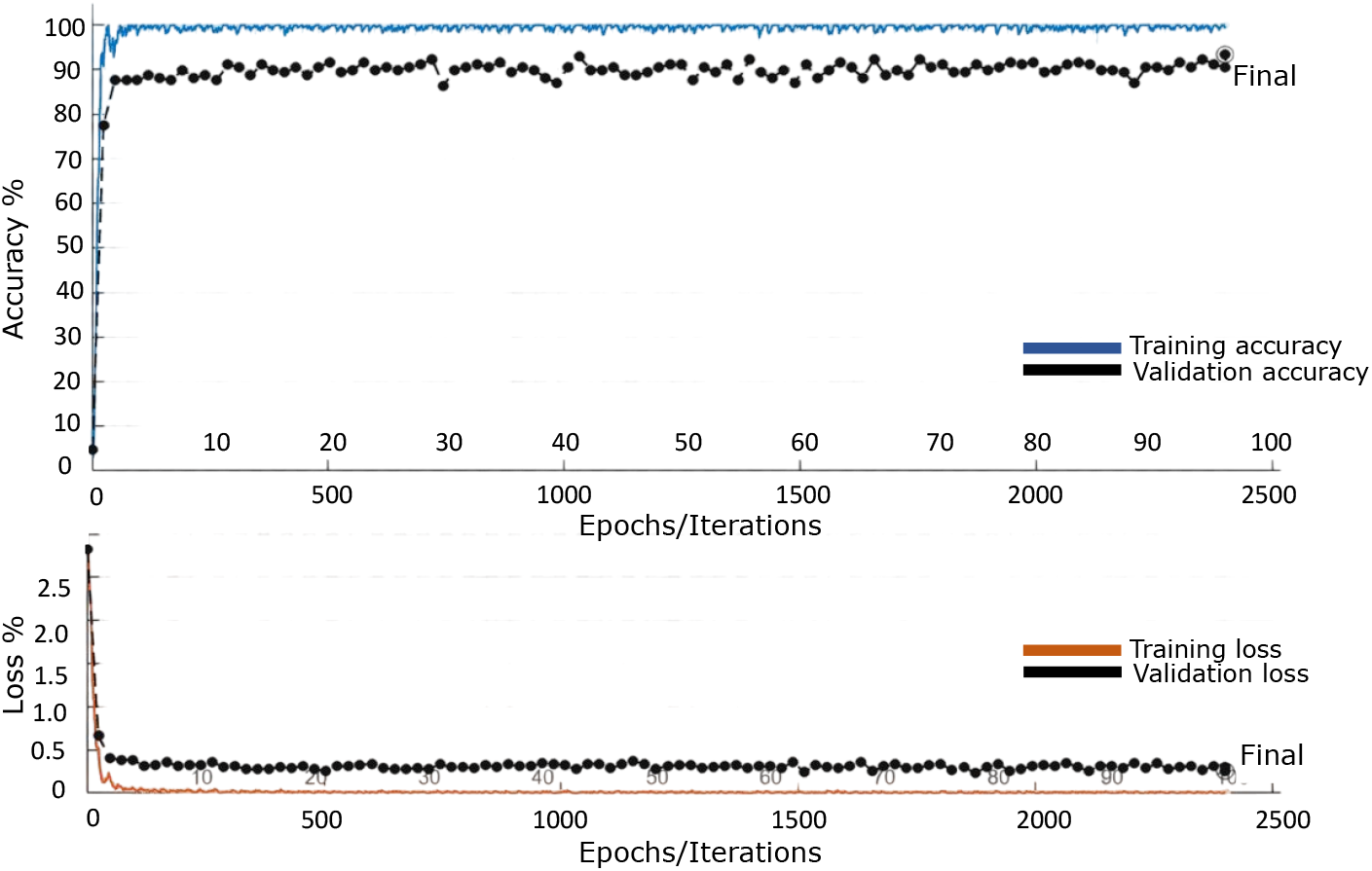
Illustrates training and validation performance of ResNet101. The model achieved an optimal training accuracy of 98% with a training loss of 0.10%, and a validation accuracy of 91% with a validation loss of 0.2%. The model was trained using a learning rate of 0.001, batch size of 32, dropout rate of 0.5, over 100 epochs and 2500 iterations.

**Figure A3.**
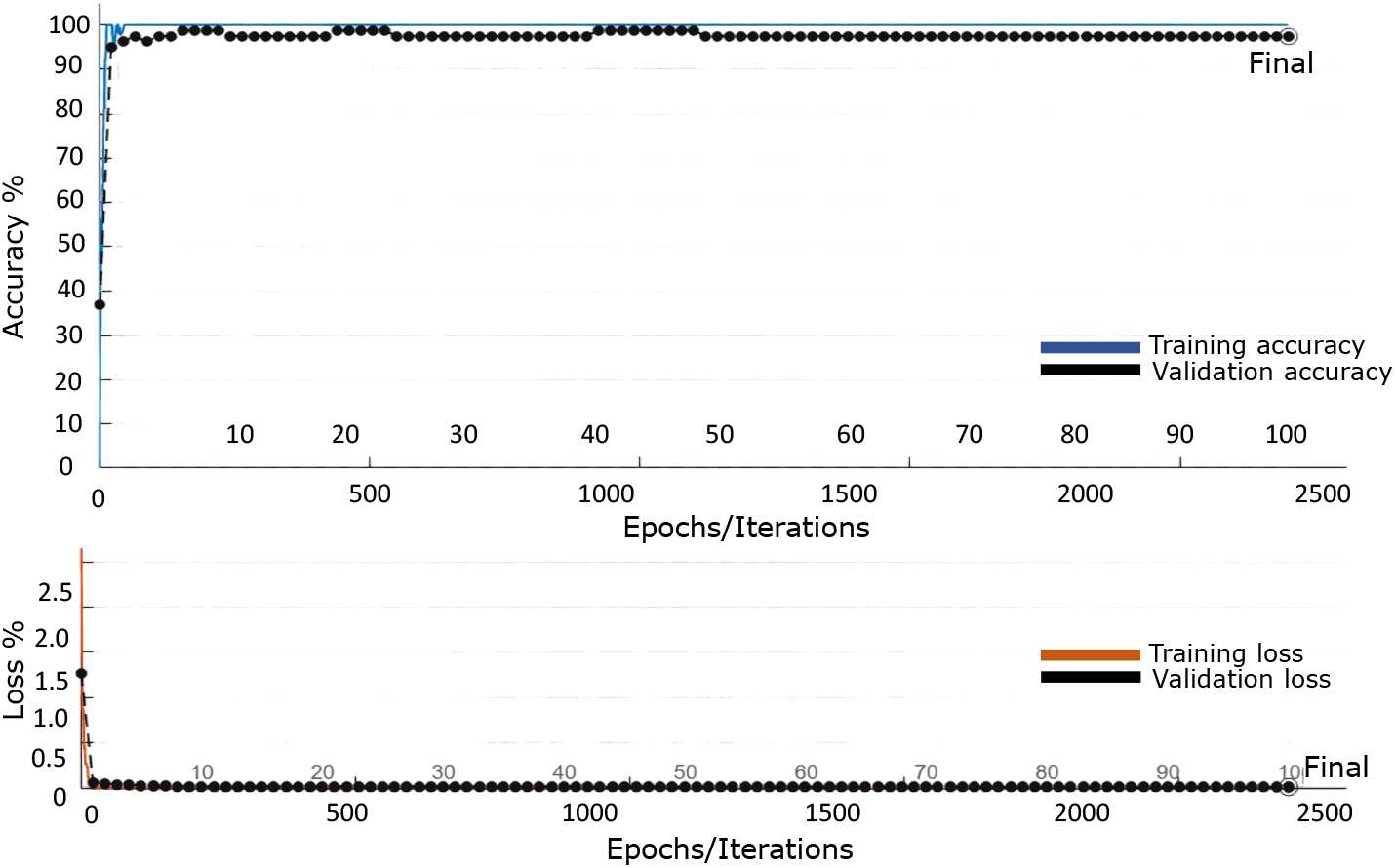
Illustrates training and validation performance of Inception-ResNetV2. The model achieved an optimal training accuracy of 100% with a training loss of 0.0001%, and a validation accuracy of 98.95% with a validation loss of 0.005%. The model was trained using a learning rate of 0.001, batch size of 32, dropout rate of 0.4, over 100 epochs and 2500 iterations.

### A.2 Displacement fields and von Mises stress distributions: Predictions vs. DIC measurements

**Figure A4.**
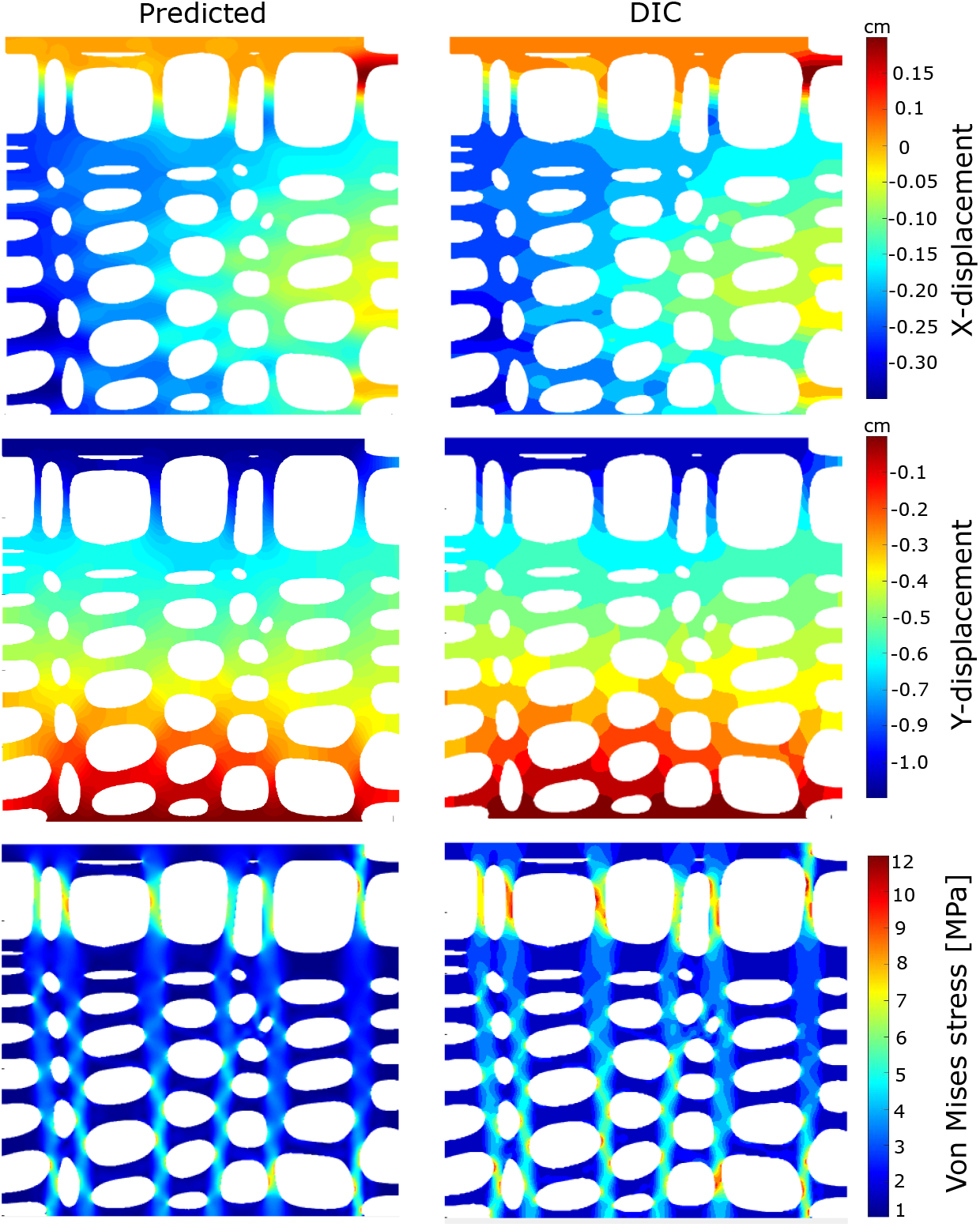
Comparison of X-displacement, Y-displacement, and von Mises stress color maps of the wood cellular structure obtained from digital image correlation (DIC) measurements and purposed CNN-based predictions

**Figure A5.**
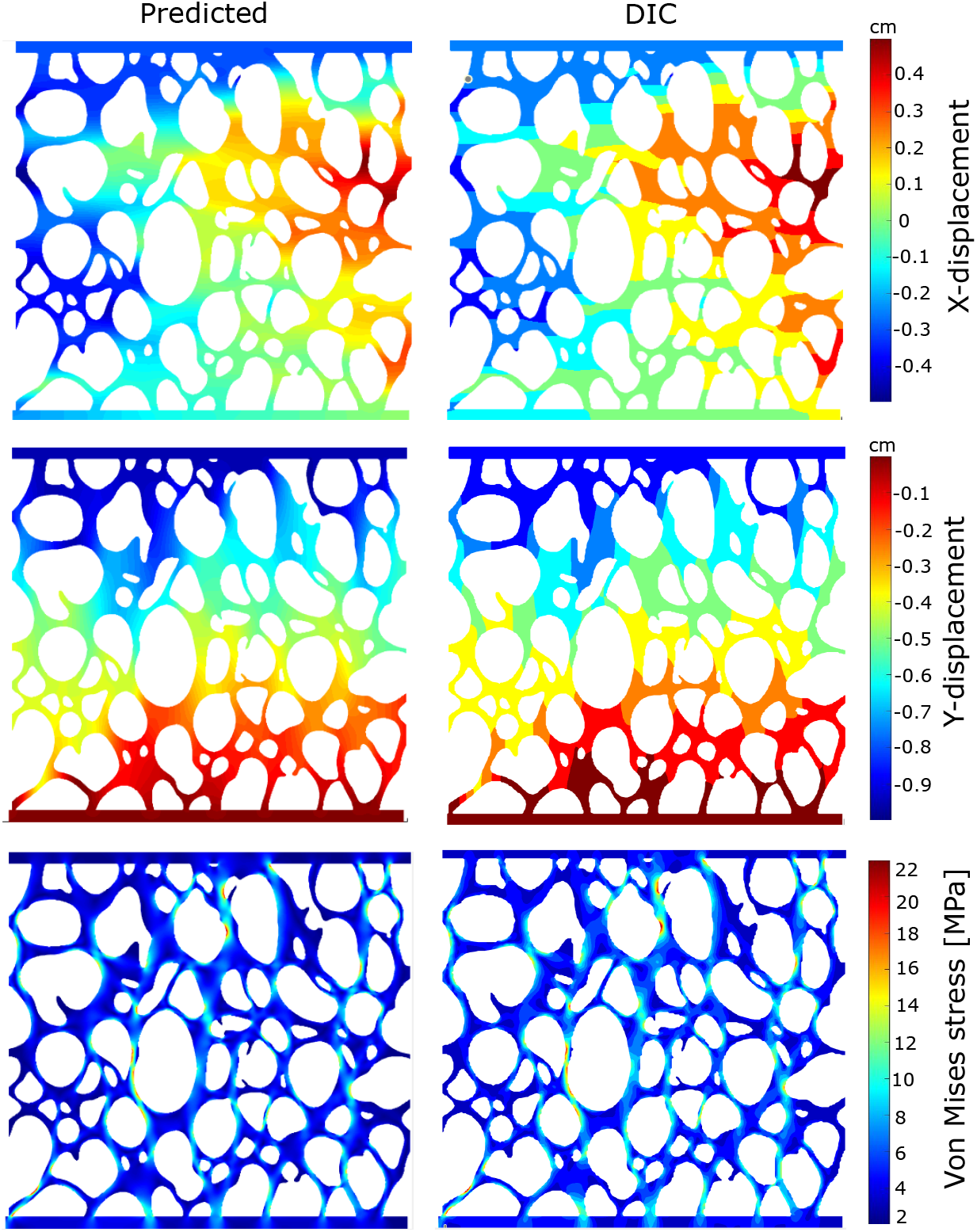
Comparison of X-displacement, Y-displacement, and von Mises stress color maps of the sponge cellular structure obtained from digital image correlation (DIC) measurements and purposed CNN-based predictions

