## Supplementary material for "AI-BioMech: Deep Learning Prediction of Mechanical Behavior in Aperiodic Biological Cellular Materials": AIBioMech User Manual

### AI-BioMech User Manual

Haleema Sadia[1],Marcelo Dias[1], Parvez Alam[1*]
[1]School of Engineering, The University of Edinburgh, Edinburgh, UK

February 20, 2026

**AI-BioMech** is a deep learning–based software framework designed to predict the mechanical response of biological cellular materials directly from 2D images. By employing state-of-the-art convolutional neural network architectures, AI-BioMech eliminates the need for the manual definition of geometries coupled to finite element anlaysis (FEA) simulation. The framework is trained on synthetic datasets that mimic biological cellular structures, while FEA-based labeling provides pixel-level annotations for semantic segmentation of stress and strain distributions. This approach allows the model to automatically extract complex spatial and hierarchical patterns, enabling accurate predictions of mechanical responses under compression. AI-BioMech incorporates transfer learning strategies using DeepLabv3 with Inception-ResNetV2 backbones to enhance prediction accuracy and generalization, even when trained on limited datasets. Model outputs are validated against experimental measurements, including Digital Image Correlation (DIC) from experimental tests, demonstrating strong agreement with physical observations. Compared to traditional methods, AI-BioMech offers significant improvements in computational speed and scalability, achieving up to 99% prediction accuracy. By integrating advanced machine learning with biomechanics, AI-BioMech provides a practical and powerful tool for rapid stress–strain estimation opens new opportunities for accelerating research in bio-inspired structures. The software nevertheless also includes an FEA simulator in both tension and compression, enabling the user to choose traditional FEA simulation when desired.

### 1 Introduction

Predicting the mechanical behavior of biological cellular materials is often a complex challenge in biomechanics and material science. This is because biological materials exhibit highly irregular geometries, making traditional modeling approaches, such as manual geometry reconstruction and finite element analysis (FEA), time consuming, computationally intensive, and prone to errors, especially when dealing with high-resolution images. To address these challenges, we present **AI-BioMech**, a deep learning–based software framework that directly predicts the mechanical response of cellular materials from high-resolution 2D images. By utilising convolutional neural networks and semantic segmentation, AI-BioMech automatically extracts complex spatial and hierarchical patterns from labeled data, enabling accurate predictions of mechanical responses under compression. The framework supports both **elastic and elastic-plastic be-**

**havior**, providing users two options to simulate realistic material responses by specifying key material properties, including **Young's modulus, Poisson's ratio, and yield strength**.

Key features of AI-BioMech include:

- **High-resolution image processing:** Handles detailed cellular geometries without loss of information.
- **Automated feature extraction:** Learns spatial and hierarchical patterns directly from FEA-labeled data.
- **Elastic and elastic-plastic behavior modeling:** Supports both linear elastic and bi-linear elastic-plastic responses with customizable material parameters.
- **Transfer learning support:** Uses DeepLabv3 with Inception-ResNetv2 backbones for improved accuracy and generalization.
- **Rapid predictions:** Offers significant speed improvements over traditional FEA simulations for fast design exploration.
- **Experimental validation:** Model outputs are benchmarked against Digital Image Correlation (DIC) measurements and physical experiments to ensure accuracy and reliability.
- **Applications in engineering and research:** Enables rapid stress–strain estimation, 2D material design exploration, optimization of cellular materials, and analysis of bio-inspired structures.
- **User-friendly GUI:** Provides an intuitive interface for easy input, parameter selection, and visualization of results.

**Please note that AI-BioMech may take a little time to start up (up to a few minutes depending on system specs). Please be patient, once you are in the GUI, it runs faster.**

### 2 Run Method

The **AI-BioMech software** can be executed using one of the following two methods, depending on user preference and system configuration.

#### 2.1 Method 1: Running AI-BioMech Using MATLAB Runtime

This method allows users to run AI-BioMech directly using the MATLAB Runtime without installing the full software package. We recommend this method as it is the fastest and lightest option for your computer.

- Install the **MATLAB Runtime (R2024a)** appropriate for your operating system (Windows, macOS, or Linux). The **AI-BioMech** application is compatible with all major operating systems; however, users must ensure that the MATLAB Runtime version installed matches their specific system architecture. MATLAB Runtime enables execution of AI-BioMech software without requiring a MATLAB license.

- MATLAB Runtime can be installed using one of the following options:

  - Download the runtime from the official MathWorks website: MathWorks MATLAB Runtime download. Select version **R2024a (v24.1)** corresponding to your operating system, as shown in Figure 1; **or**
  - Use the MATLAB Runtime setup folder provided with the AI-BioMech software (for Windows users). Extract the folder to a preferred location and run the **setup** file to install the runtime.

- After successful installation of the MATLAB Runtime, locate the executable file `AIBioMech.exe`.

- Double-click `AIBioMech.exe` to launch the software. The application opens the **AI-BioMech graphical user interface (GUI)**, enabling the straightforward input of high-resolution images and material parameters.

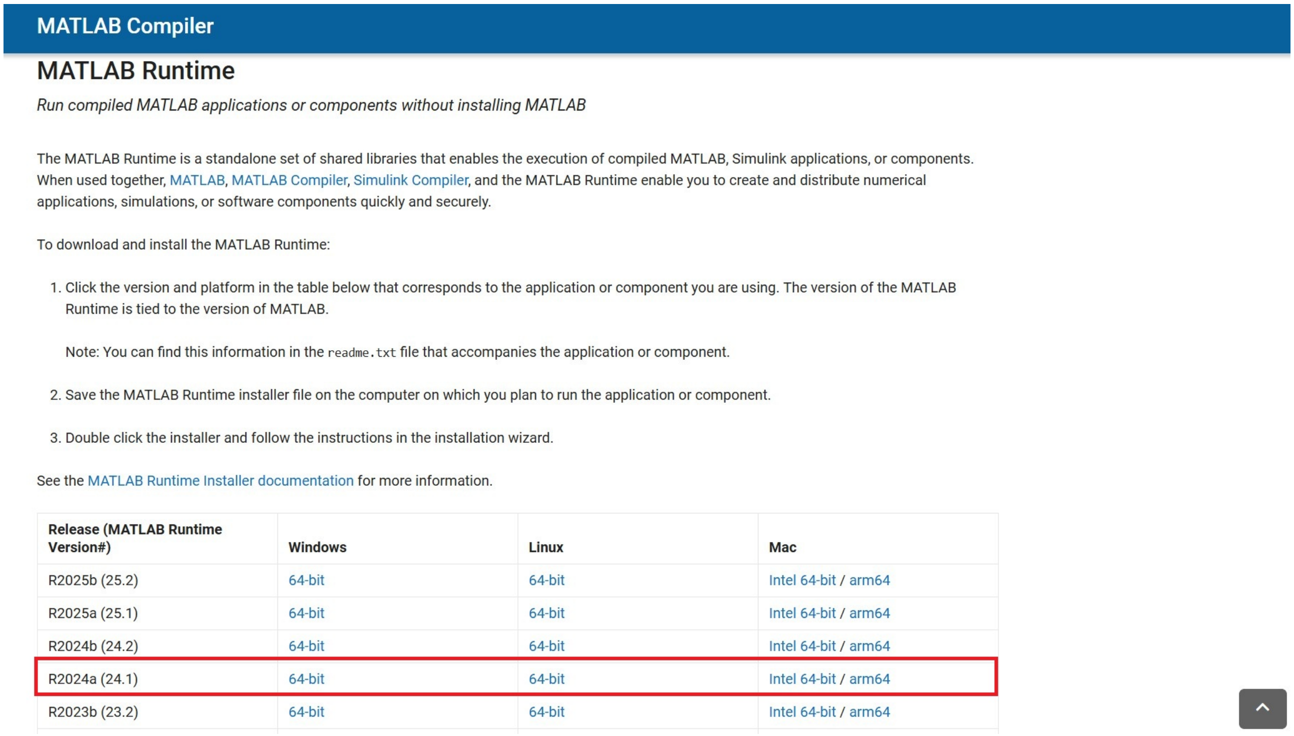


Figure 1: MATLAB Runtime setup (version 24.1) compatible with AI-BioMech software

#### 2.2 Method 2: Running AI-BioMech Using the Installer Package

This method installs AI-BioMech as a standalone application using the provided installer.

- **Download the AIBioMech.zip package**: Download the complete deployment package. This package includes the MATLAB Runtime, enabling the software to run independently. The **AIBioMech.zip** file is approximately 3.91 GB in size, and download time depends on internet speed and system performance.
- **Extract AIBioMech.zip**: Extract the contents of the zip file to a directory of your choice. It is highly recommended that you use a location that does not require special permissions.
- The extracted **AIBioMech** folder contains the following file:
  1. **MyAppInstaller_mcr**
- Extract the **MyAppInstaller_mcr** folder and run the **MyAppInstaller_mcr** setup file. Upon execution, the **AIBioMech Installer** window will open, as shown in Figure 2.

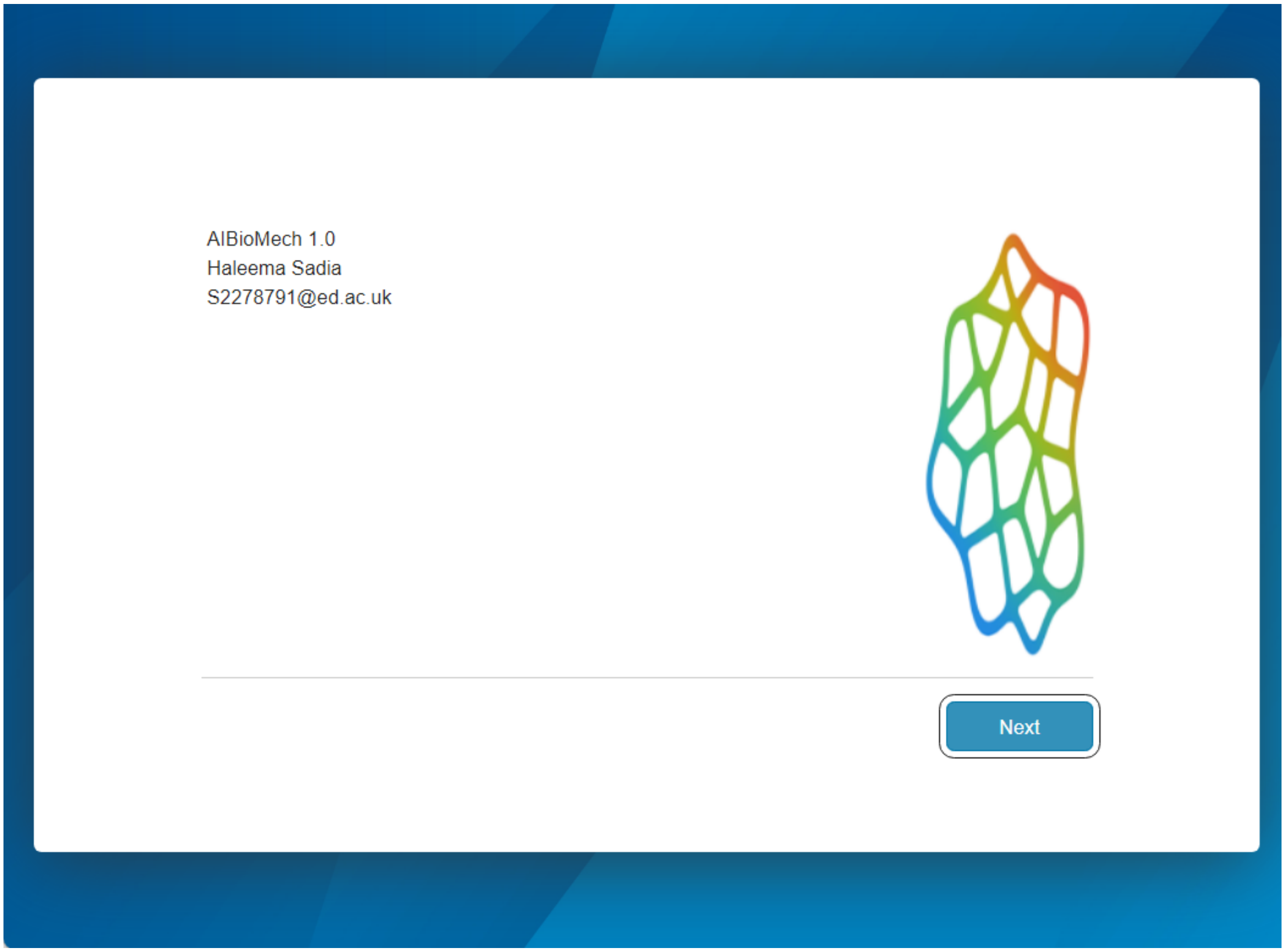


Figure 2: AIBioMech installer

- Click the **Next** button to proceed. The Installation Folder window will appear (Figure 3). Select the desired installation directory (e.g., `Program Files`) and choose whether to create a desktop shortcut.

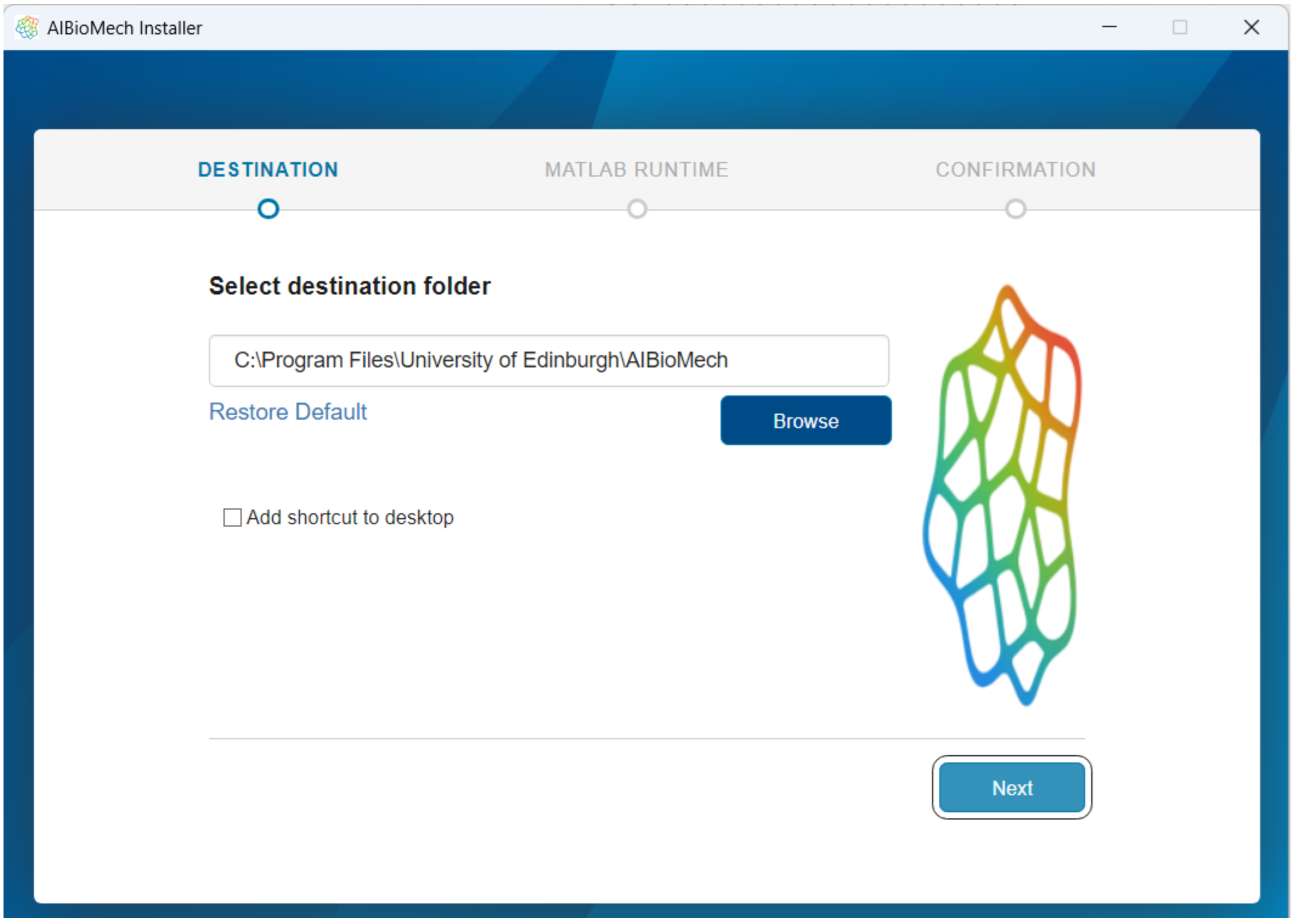


Figure 3: AIBioMech installation destination folder path

- Click **Next** to open the confirmation window (Figure 4).

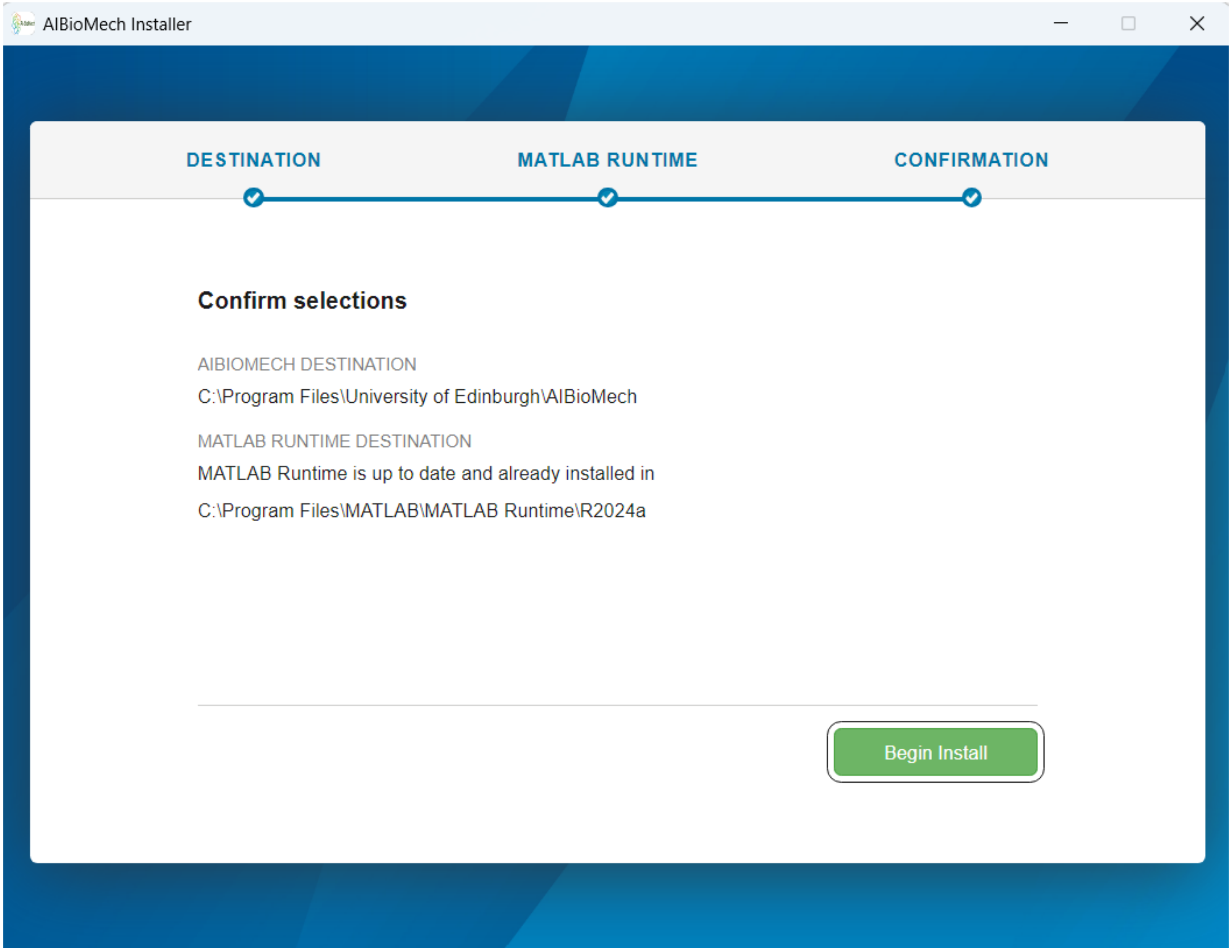


Figure 4: AI-BioMech installation confirmation

- Press the **Begin Install** button to begin the installation. The process may take several minutes.
- After installation is complete, the completion window will appear (Figure 5). Click the **Close** button to exit the installer.

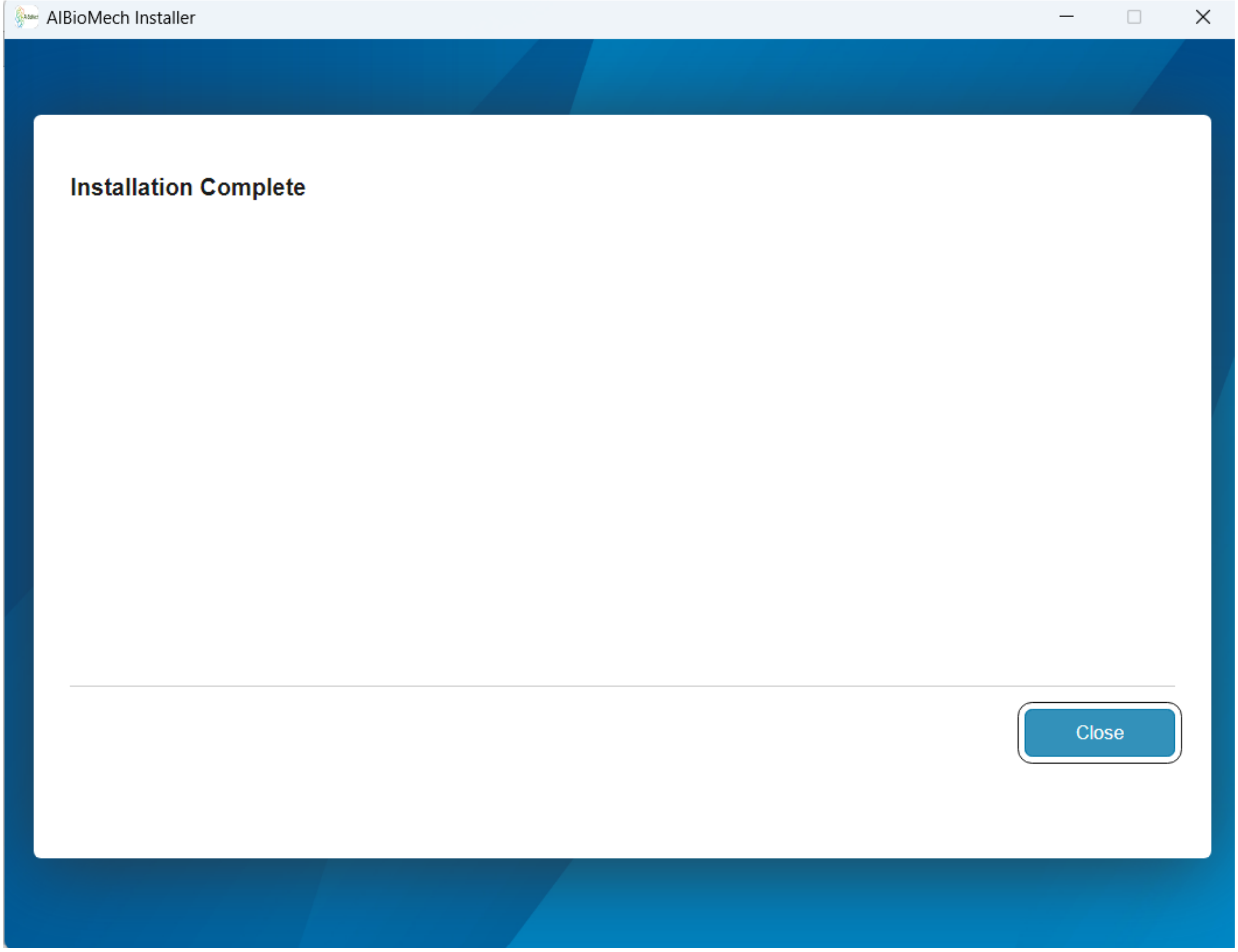


Figure 5: AIBioMech installation completion

- **Run AIBioMech**: Launch the software by double-clicking the **AIBioMech** desktop shortcut or by opening the executable from the installation directory.
- **readme**: The **readme** file included with the package provides essential information regarding system requirements, usage instructions, and deployment details.

### 3 AI-BioMech Graphical User Interface (GUI)

After Double-click `AIBioMech.exe` to launch the software, the Graphical User Interface (GUI) will appear, Figure 6.

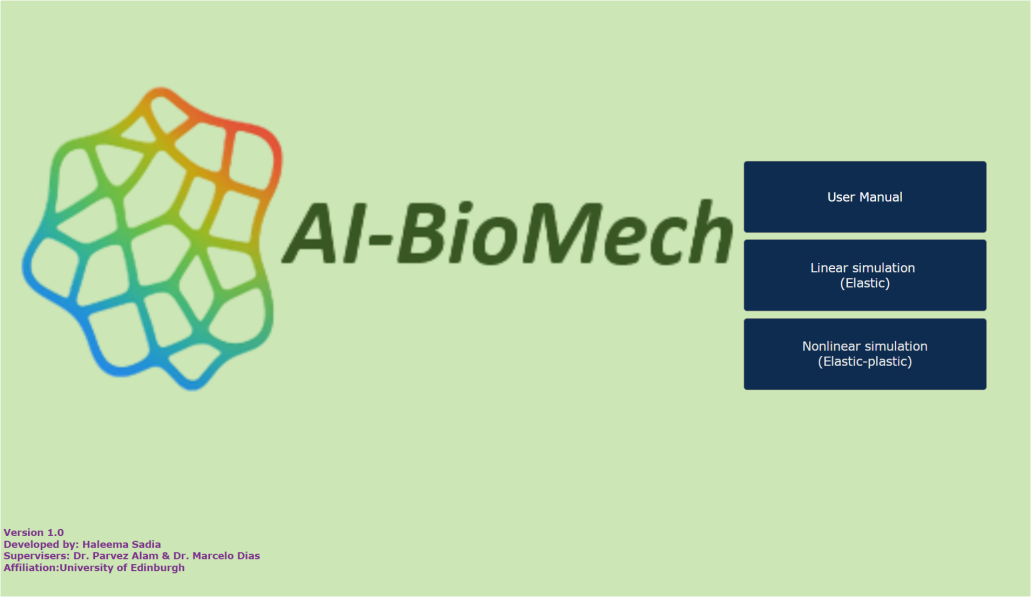


Figure 6: Main workspace: Graphical user interface (GUI) of AI-BioMech

The main interface provides three buttons to guide the user through the software workflow:

- **User Manual:** Press this button for the user manual, includes instructions for software run and usage.
- **Linear simulation (Elastic):** Press this button to run simulations assuming linear elastic material behavior.
- **Nonlinear simulation (Elastic–plastic):** Press this button to run simulations for materials exhibiting both elastic and plastic behavior.

Both simulation panels follow the same workflow for image loading, region of interest selection, scale setting, and material properties input.

The primary difference lies in how material behavior is modeled:

- **Linear Simulation Panel**: Assumes linear elastic material behavior, where stress is linearly proportional to strain. Users only need to enter the **Young's modulus** and **Poisson's ratio** for the material, along with displacement values. The resulting stress and strain distributions are computed using linear elasticity theory.
- **Nonlinear Simulation Panel**: Accounts for material nonlinearity by requiring the user to additionally specify the **yield stress** in the material properties field. This enables the software to simulate plastic deformation beyond the elastic limit. All other steps, including **Young's modulus** and **Poisson's ratio** for the material, along with displacement values, remain the same as in the linear simulation workflow.

#### 3.1 Linear simulation (Elastic)

Pressing the **Linear simulation (Elastic)** button opens a new window that provides the GUI for linear simulation, as shown in Figure 7.

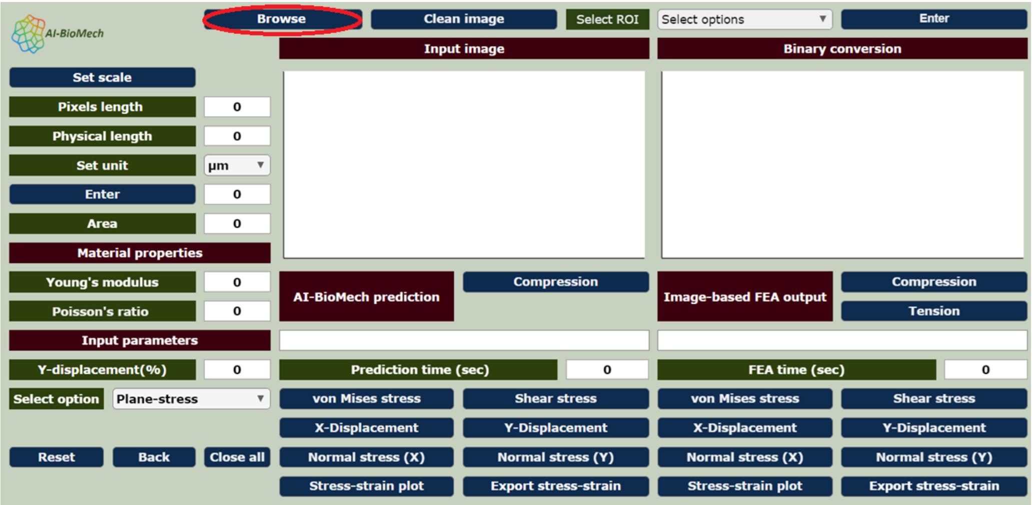


Figure 7: AI-BioMech graphical user interface (GUI) for linear elastic simulations

Press the **Browse** button to select a high-resolution 2D image of the biological cellular structure (supported formats: .png, .tiff, or .jpg). Once the image is loaded, the GUI displays:

- The original RGB image on the first axis, labeled as **Input Image**.
- The corresponding binary image on the second axis, labeled as **Binary Conversion** (As shown in Figure 8).

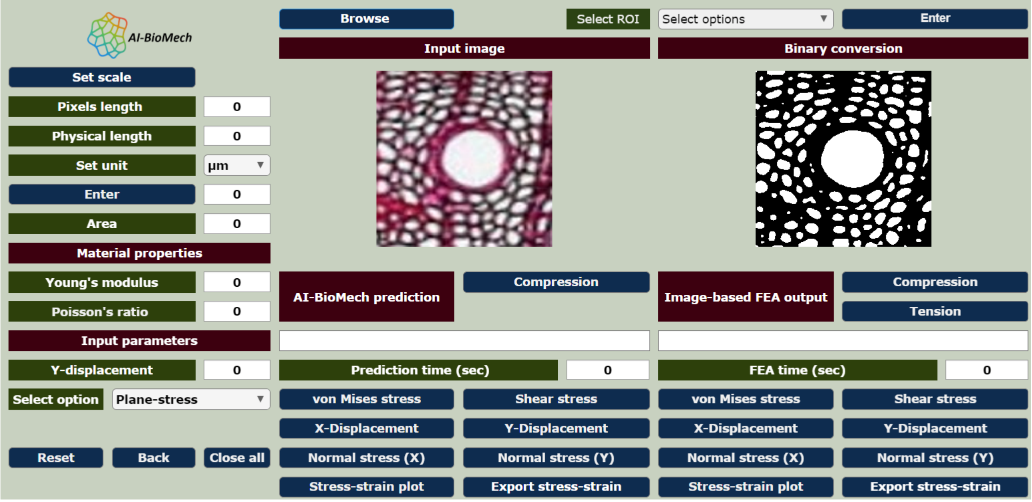


Figure 8: Graphical user interface (GUI) of AI-BioMech linear simulation: Browse button

When an image is noisy or contains irregular pixels, these imperfections can negatively affect segmentation accuracy, mesh generation, and simulation results. By using the **Clear Image** button, users can interactively clean and refine the image to improve overall quality as shown in Figure 9.

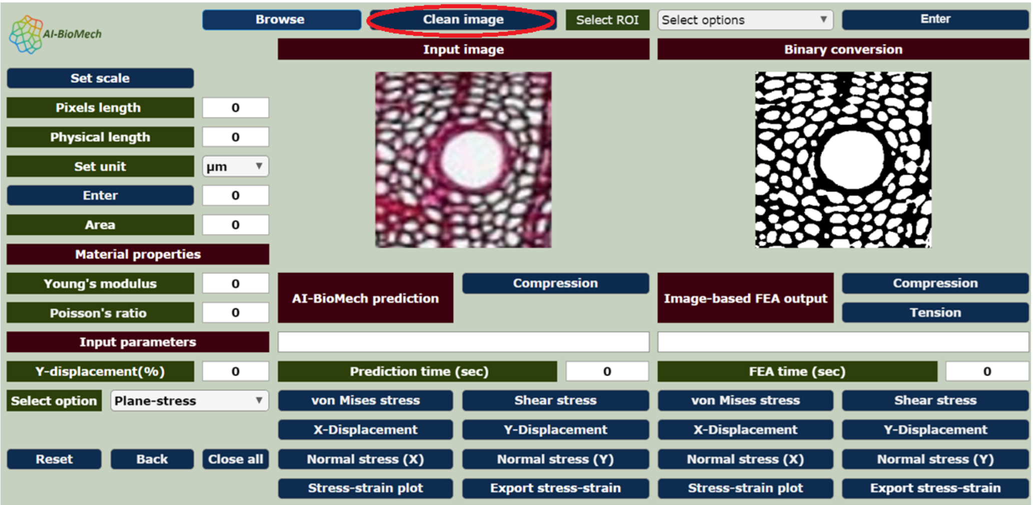


Figure 9: AI-BioMech graphical user interface (GUI) for linear elastic simulations: Clear image

When the user clicks the **Clear Image** button, a new window opens that provides three image editing options (See Figure 10): **Erosion**, **Dilation**, and **Eraser**. It shows the total number of pixels (white & black) The user can adjust the parameters of each operation using the corresponding sliders.

- **Erosion and Dilation:** These operations are used to refine image features by removing noise or enhancing structural continuity. The intensity of each operation can be controlled through the slider.

- **Eraser:** The eraser allows interactive removal of unwanted pixels from the image. If the user clicks on a **black region**, black pixels are erased. If the user clicks on a **white region**, white pixels are erased. The size of the eraser can be adjusted using the eraser size slider .

After setting the desired parameters, the user must **double-click the "Save and Exit" button** to confirm and apply the changes. Once saved, the processed image is used for all subsequent steps, including prediction and finite element analysis (FEA) simulation.

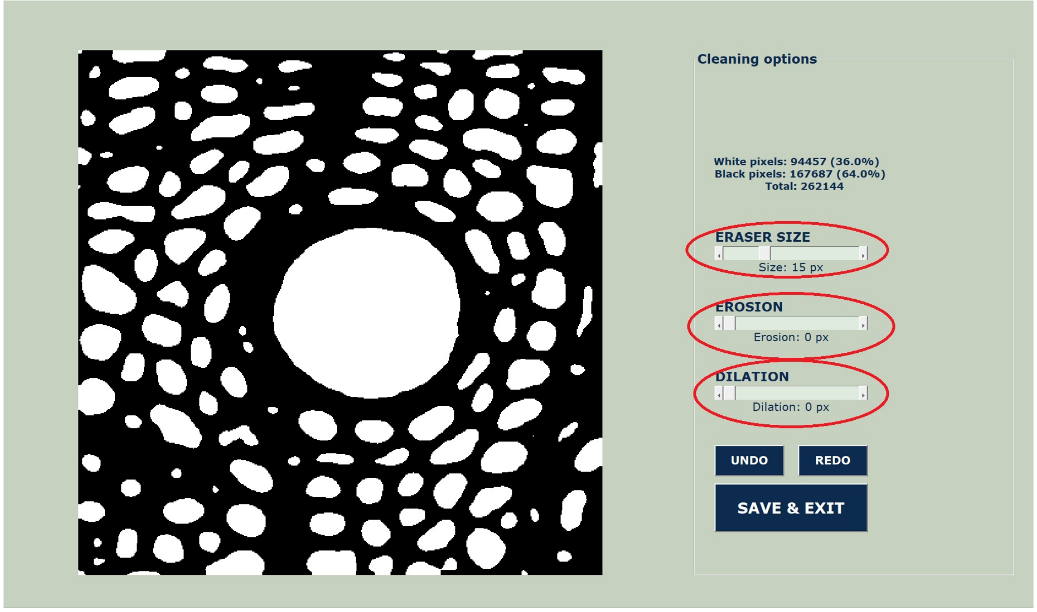


Figure 10: AI-BioMech graphical user interface (GUI) for linear elastic simulations: Clear Image options

After browsing the image, select the **region of interest (ROI)** corresponding to the cellular biological structure. Choose the appropriate ROI type from the drop-down menu and press **Enter** to proceed as shown in Figure 11.

**Important Note: The input image must be high-resolution and square in shape. If the input image is not square, it will be automatically resized to $1024 \times 1024$, which may cause image distortion, squeezing, or unintended alteration of the original geometry. The region of interest (ROI) should aim to minimize any floating, disconnected, or free boundaries. Low image resolution or poor image quality may result in mesh generation errors during image-based finite element analysis (FEA).**

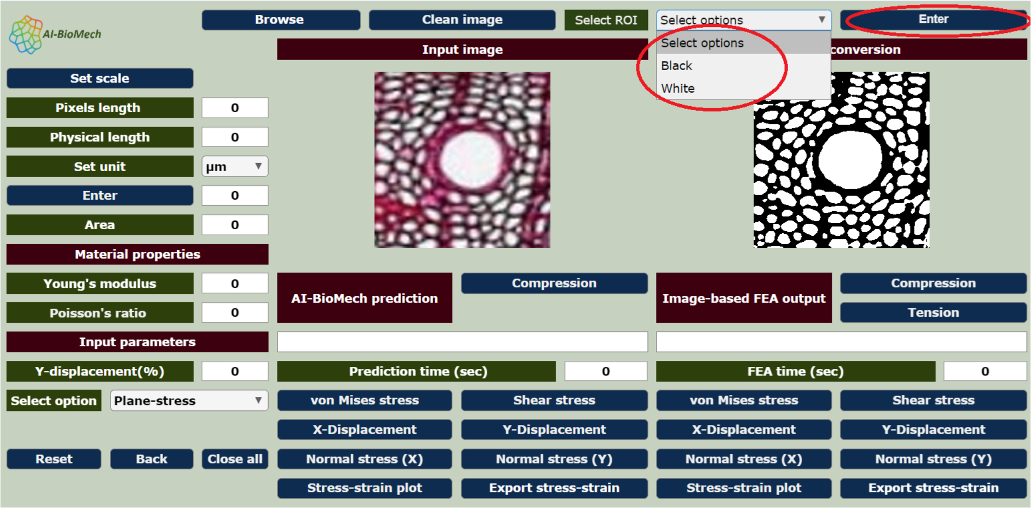


Figure 11: AI-BioMech graphical user interface (GUI) for linear elastic simulations: Select Region of interest (ROI)

The Region of Interest (ROI) refers to the specific area of the image that contains the object or structure to be analyzed. Only this region is used for processing. The ROI should clearly include the complete geometry of interest without any unnecessary background, noise, or isolated regions. After selecting the Region of Interest (ROI) from the loaded cellular biological image, the user must press the **Enter** button at the top right of the GUI, as seen in the figure. Use then, the **Set Scale** button to calibrate the image according to a known physical dimension as shown in Figure 12. This step allows the software to convert pixel-based measurements into meaningful physical units. After double-clicking on the **Pixel Length** measurement, the value is automatically displayed in the **Pixel Length** field. The user should enter the corresponding physical length of the selected reference in the provided **Physical Length** field and then select the appropriate SI unit (such as micrometer, millimeter, centimeter or inch) from the units drop-down menu. Once the physical length and unit are specified, pressing the **Enter** button below the **Set Unit** button, applies are scaling to the image. After the scale is successfully set, the software computes and displays the area and length measurements of the selected region using the defined physical scale.

**Note: If the user does not set the scale, all measurements and subsequent simulations are performed using the default pixel-based scale, and the results are reported in pixel units.**

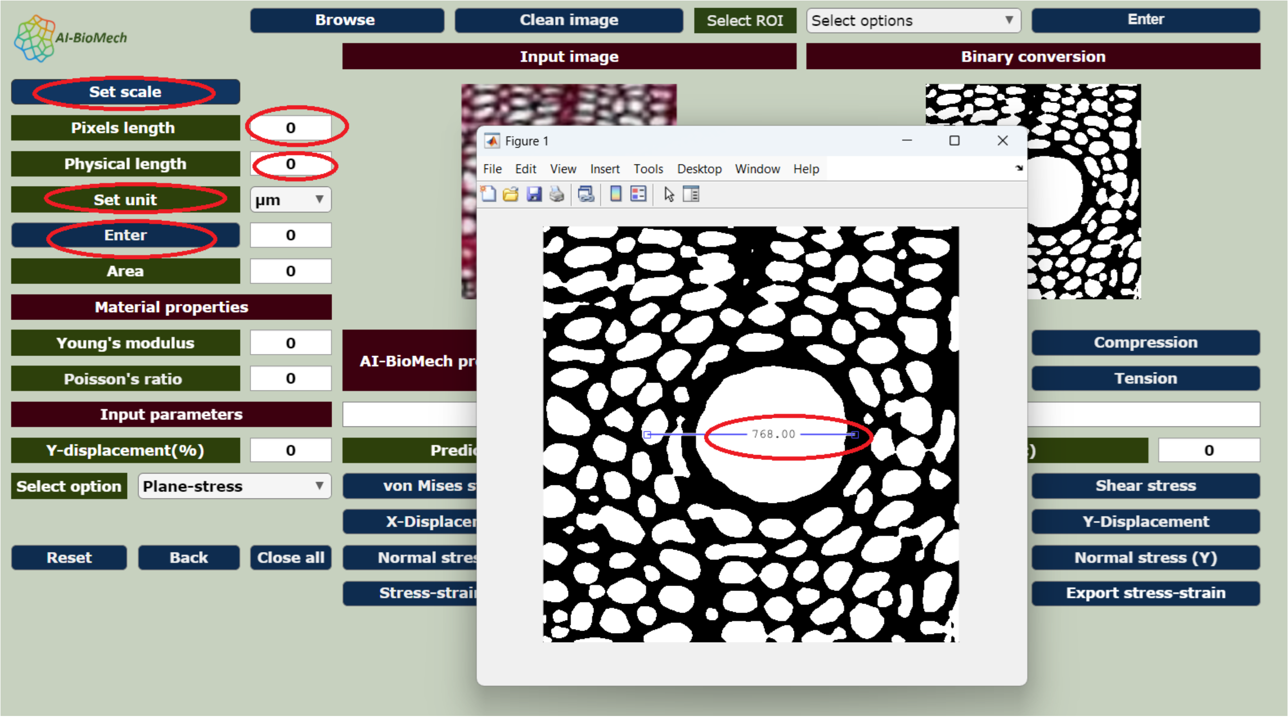


Figure 12: AI-BioMech graphical user interface (GUI) for linear elastic simulations: Set scale

After the scale has been successfully set, the user is required to define the **Material properties** for the analysis. This includes entering the **Young's modulus** and **Poisson's ratio** in their respective input fields. These parameters describe the mechanical behavior of the selected material and are essential for obtaining accurate simulation results. Under **Input parameters**, the user must specify the **Y-direction displacement** to define the applied boundary condition. The displacement value must be entered as a **percentage of the total specimen length**. For example, a value of **0.1** corresponds to a displacement equal to **0.1%** of the total specimen length. Users should enter only **positive values**; the software internally interprets the displacement as either **compressive or tensile loading** based on the selected analysis mode. The user should then select the appropriate analysis type from the **drop-down menu**, choosing between **plane stress** and **plane strain**, depending on the type of 2D simulation required, Figure 13.

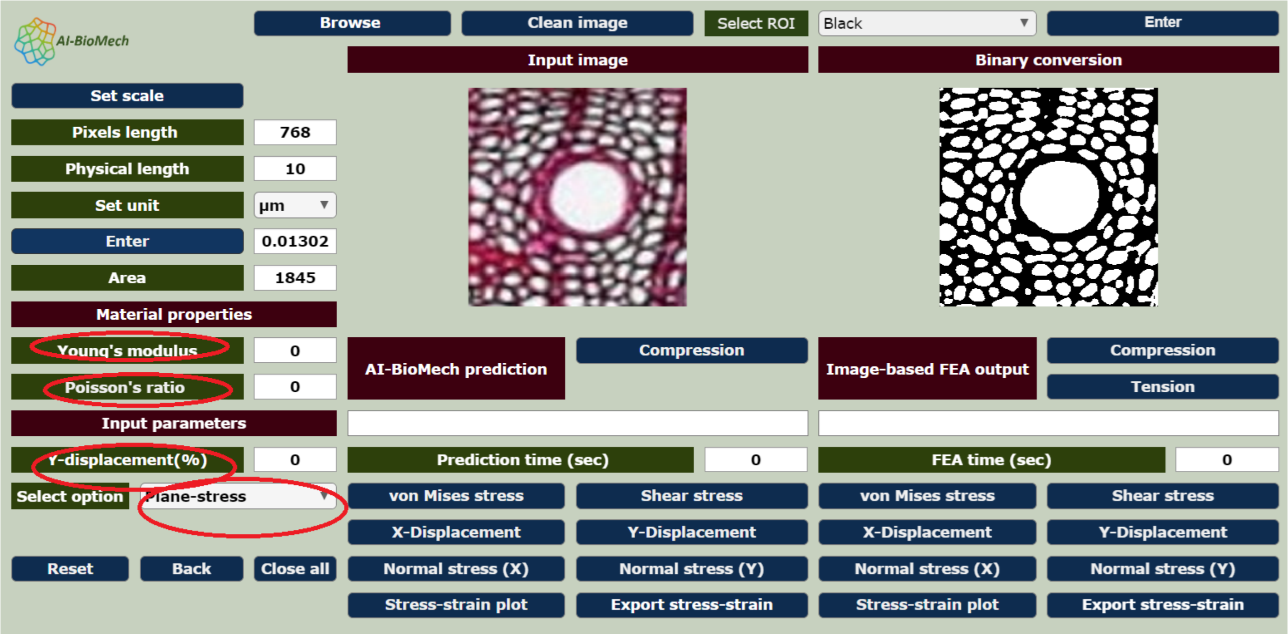


Figure 13: AI-BioMech graphical user interface (GUI) for linear elastic simulations: Set material properties and input parameters

After defining the material properties and analysis type, the user is directed to the two separate panels on the GUI: **AI-BioMech Prediction** and **Image-Based FEA output** as shown in Figure 14. The Image-Based FEA panel is intended to generate **ground-truth simulation results** directly from the image-derived geometry and physical parameters. In contrast, the AI-BioMech Prediction panel provides **predicted outputs** generated using a trained artificial intelligence model. Users can evaluate and validate the accuracy of the AI-based predictions by comparing them with the results obtained from the image-based FEA simulations. To obtain the AI-based prediction, the user should press the **Compression** button located in the AI-BioMech Prediction panel. Similarly, to perform the image-based FEA simulation, the user must press the **Compression** button in the simulation section of the Image-Based FEA panel.

**Important Note: In the current version of the software, the AI model is trained only for compressive loading. If the user wishes to perform a tensile simulation, this can be carried out using the image-based finite element analysis (FEA) panel, which supports both compression and tension cases.**

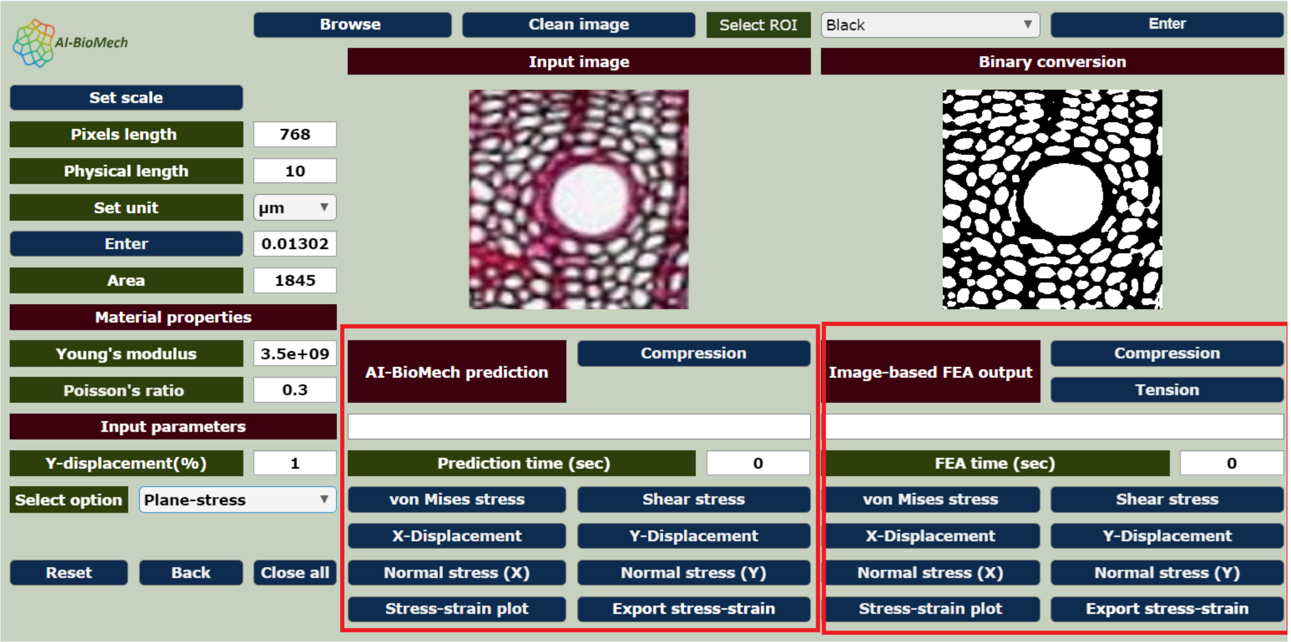


Figure 14: AI-BioMech graphical user interface (GUI) for linear elastic simulations: AI-BioMech prediction panel and Image-based FEA simulation panel

After the simulation is complete, the software displays the **maximum linear stress** and **maximum linear strain** values in the corresponding text fields for both the AI-based prediction and the image-based FEA simulation. In addition, the **simulation time** required for each method is also reported. These outputs enable users to directly compare the computational efficiency and predictive accuracy of the **AI-BioMech** framework with the conventional image-based FEA approach. By analyzing differences in simulation time and stress–strain results, users can assess the performance, efficiency, and reliability of the AI-based biomechanical predictions. The software provides several interactive buttons to aid in the visualization of simulation results, Figure 15. These **buttons** include:

- **von Mises stress**: Displays the **von Mises stress distribution** over the selected region using a color map representation, allowing visualization of stress concentration zones.
- **Shear stress**: Generates the **shear stress distribution map**, highlighting regions subjected to shear loading within the structure.
- **X-Displacement**: Visualizes the **displacement field in the X-direction** using a color map to illustrate horizontal deformation.
- **Y-Displacement**: Displays the **displacement field in the Y-direction**, representing vertical deformation across the structure.
- **Normal stress (X)**: Shows the **normal stress distribution in the X-direction**, enabling assessment of compressive stresses acting horizontally.
- **Normal stress (Y)**: Displays the **normal stress distribution in the Y-direction**, providing insight into vertical stress behavior.

- **Stress–strain plot**: Plots the **stress–strain relationship** derived from the prediction/simulation results, facilitating mechanical behavior analysis and comparison between AI-based prediction and image-based FEA.

- **Export stress–strain**: Allows the user to **export stress–strain data** for further analysis, reporting, or validation using external tools.

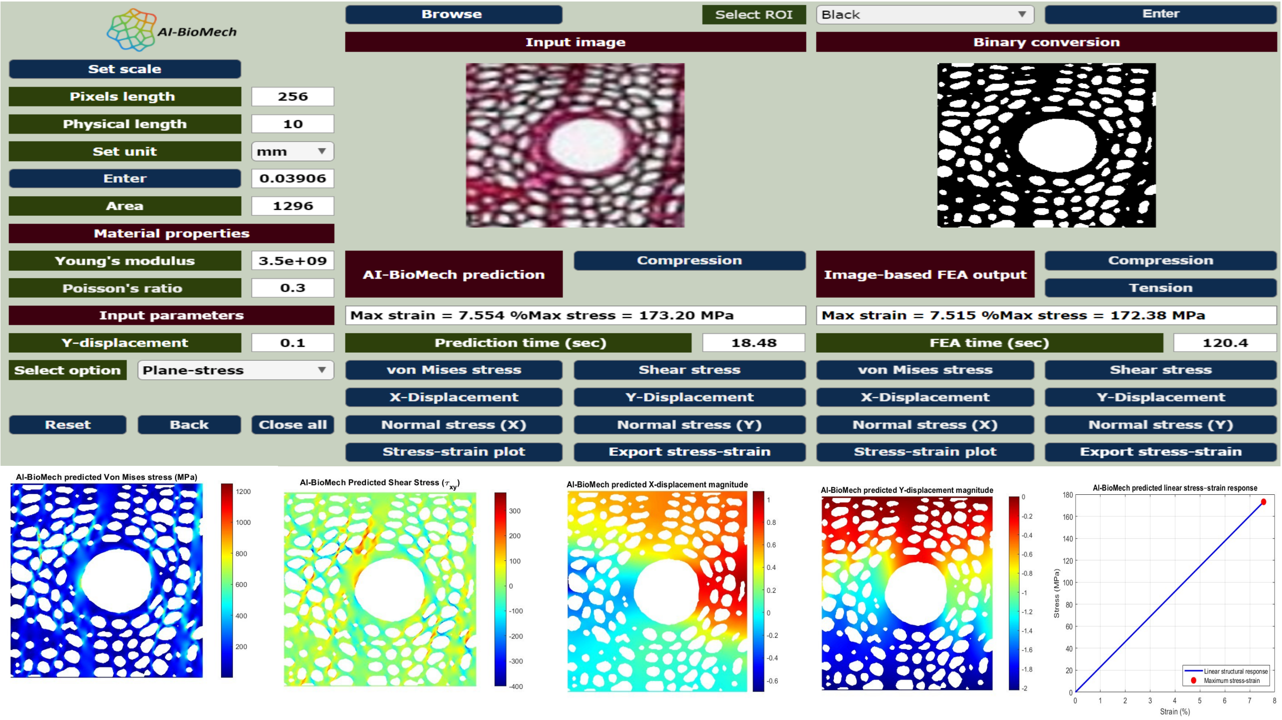


Figure 15: AI-BioMech graphical user interface (GUI) for linear elastic simulations: Output Visualization

- **Close figures**: Closes all currently open figure windows or plots generated by the software, helping to declutter the workspace.

- **Reset**: Clears all inputs, selections, and displayed images in the app, returning the interface to its default initial state.

- **Back**: Returns the user to the previous main workspace, restarting the entire workflow.

**Note: if you wish to load and simulate a new image using the 'Browse' button, you must select the ROI and press 'Enter', else the previous image will be simulated as opposed to the new uploaded image. Inputs on the left of the GUI can be changed or kept the same depending on the simulation requirements, but the ROI must be chosen and 'Enter' pressed.**

#### 3.2 Nonlinear simulation (Elastic-plastic)

**Nonlinear simulations** in AI-BioMech are based on bilinear elastic-plastic models, accounting for material nonlinearity through the addition of an additional input, the **yield strength**, in the **Material properties** field as shown in Figure 16. The software uses this value to simulate plastic deformation beyond the elastic limit. All other steps, including ROI selection, scale calibration, material properties, and displacement input, remain the same as in the linear simulation workflow. The nonlinear panel therefore provides more bi-linear predictions for materials that undergo plastic deformation under compressive loading (AI BioMech) or compressive and tensile loading (Image-Based FEA output, only).

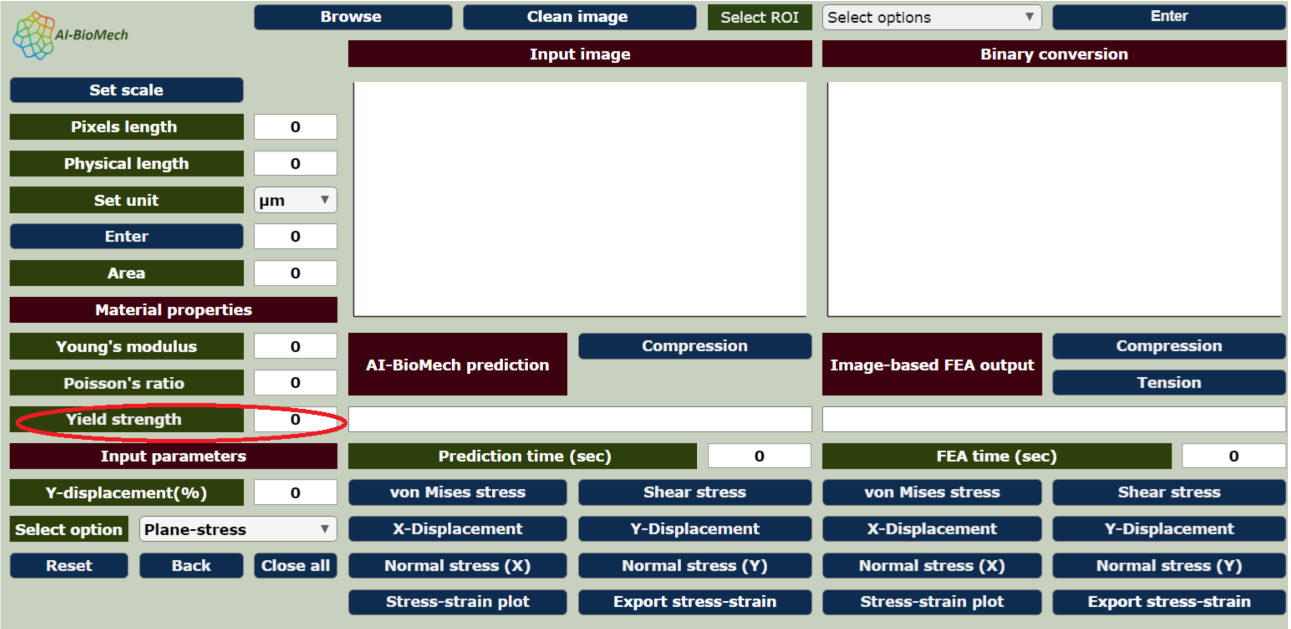


Figure 16: AI-BioMech graphical user interface (GUI) for linear elastic simulations: nonlinear simulation

Nonlinear simulation displays a **bilinear plot** representing the material response. The plot illustrates two linear regions: the initial elastic behavior followed by a secondary linear region corresponding to plastic or post-yield deformation (see Figure 17). If the user wishes to predict the **nonlinear behavior** of the cellular structure, including plastic deformation beyond the elastic limit, this simulation panel must be used. **NOTE: elastic-plastic behavior is only available through the stress-strain plot button in each of AI-BioMech prediction and Image-based FEA output. The colors maps are currently based on linear elastic simulations.**

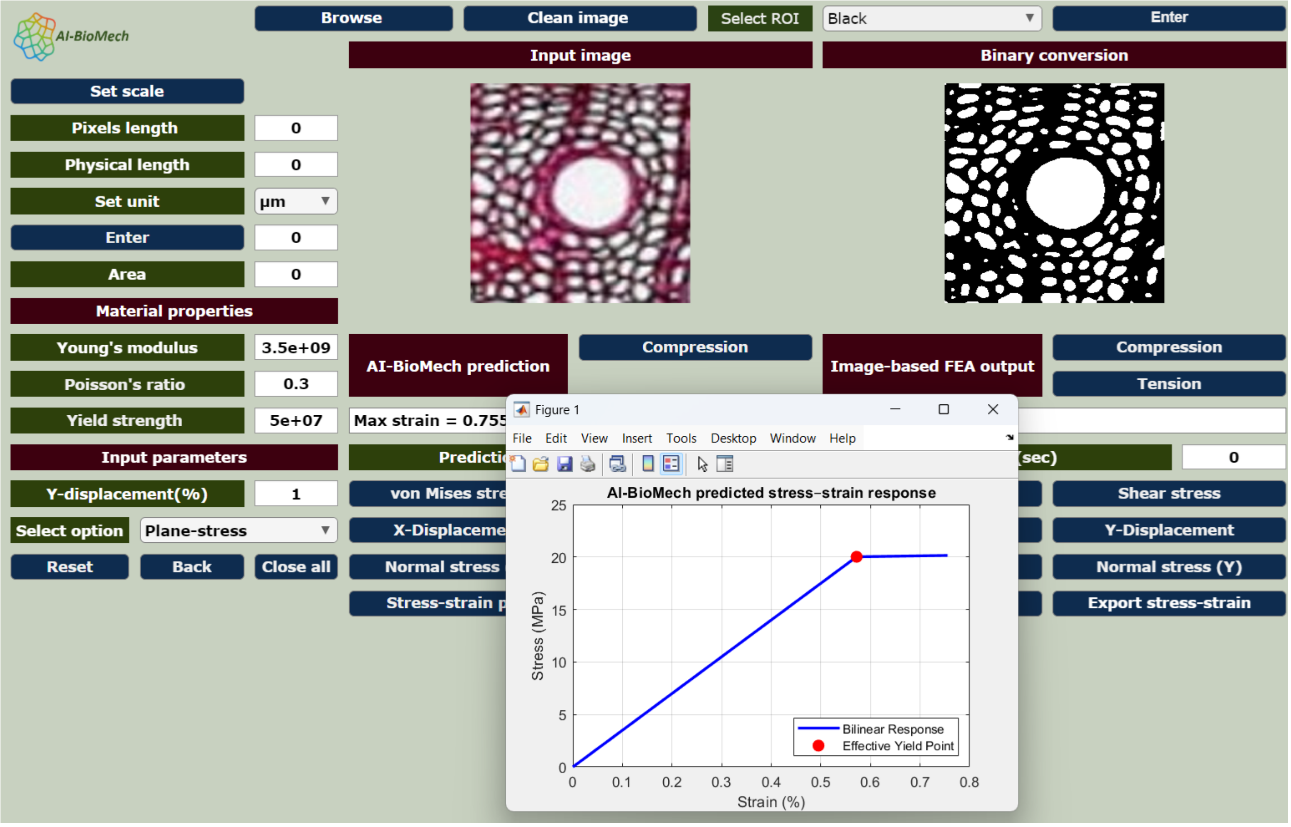


Figure 17: AI-BioMech graphical user interface (GUI) for linear elastic simulations: bilinear plot

### Constraints, limitations and additional considerations

AI-BioMech predicts the stress–strain behavior of cellular materials under prescribed boundary and loading conditions. While high predictive accuracy has been demonstrated within the validated regime, users should be aware of the following constraints and limitations.

#### 3.3 Boundary conditions and loading assumptions

All predictions are generated assuming a fixed (Dirichlet) boundary condition applied along the bottom edge of the model and a displacement applied to the top edge. These boundary conditions are embedded in the training and validation datasets and therefore directly influence the learned response. As such, the software is not intended for the simulation of non-cellular material blocks, as the effects of the Dirichlet boundary may not be ideal for non-cellular materials. Users are therefore also advised take into consideration that any predictions made are not independent of the specific boundary conditions used in this software.

#### 3.4 Material representation and plasticity modeling

AI-BioMech is trained on linear elastic simulations. The predicted color maps are generated form these simulations and the color maps therefore represent elastic stress distributions at each load increment. The stress–strain curve output from the elastic-plastic simulation is different to the color map, as it is computed by integrating a bilinear elastic–plastic constitutive model with isotropic yielding criteria, selected for consistency with available experimental data and Digital Image Correlation (DIC) measurements. Consequently, while the spatial field predictions (stress color maps) are based on linear elastic simulations, the nonlinear response is captured by the stress–strain curves through the adopted elastic–plastic formulation.

#### 3.5 Cellular size and scale effects

The AI-BioMech model is trained on $5 \times 5$ unit-cell configurations. Although satisfactory predictive performance has been observed for larger domains (up to approximately $20 \times 20$ cells), prediction accuracy progressively degrades as the cellular size extends beyond the training regime. Users should therefore be aware that very large cellular structures may fall outside the reliable operating range of the predictor. For cellular solids with larger cell-counts, we strongly recommended that AI-based predictions be validated against image-based finite element analysis (FEA) simulations as this will provide direct evidence as regards the reliability of AI predictions far beyond the training regimen. The AI tool is intended to complement, rather than replace, numerical modeling, particularly when extrapolating beyond the validated training domain.

#### 3.6 General applicability

AI-Biomech is a predictive aid within a clearly defined modeling framework. Predictions should not be interpreted as universally applicable under all boundary conditions. The transparent reporting of assumptions, boundary conditions, and model limitations is essential when using results generated by this tool.
